# Notch signaling via Hey1 and Id2b regulates Müller glia’s regenerative response to retinal injury

**DOI:** 10.1101/2021.03.11.435053

**Authors:** Aresh Sahu, Sulochana Devi, Jonathan Jui, Daniel Goldman

## Abstract

Zebrafish Müller glia (MG) respond to retinal injury by suppressing Notch signaling and producing progenitors for retinal repair. A certain threshold of injury-derived signal must be exceeded in order to engage MG in a regenerative response (MG’s injury-response threshold). Pan-retinal Notch inhibition expands the zone of injury-responsive MG at the site of focal injury, suggesting that Notch signaling regulates MG’s injury-response threshold. We found that Notch signaling enhanced chromatin accessibility and gene expression at a subset of regeneration-associated genes in the uninjured retina. Two Notch effector genes, *hey1* and *id2b*, were identified that reflect bifurcation of the Notch signaling pathway, and differentially regulate MG’s injury-response threshold and proliferation of MG-derived progenitors. Furthermore, Notch signaling component gene repression in the injured retina suggests a role for Dll4, Dlb, and Notch3 in regulating Notch signaling in MG and epistasis experiments confirm that the Dll4/Dlb-Notch3-Hey1/Id2b signaling pathway regulates MG’s injury-response threshold and proliferation.

## Introduction

Although the mammalian and zebrafish retina share structure and function, only zebrafish are able to regenerate a damaged retina. Key to this regenerative response are Müller glia (MG) that normally contribute to retinal structure and homeostasis (Goldman 2014, Lahne et al 2020, Lenkowski & Raymond 2014, MacDonald et al 2015, Reichenbach & Bringmann 2013, Wan & Goldman 2016). MG are the only retinal cells that exhibit a radial glial morphology with processes extending longitudinally and laterally that allow MG to interact with and sample their environment.

A needle poke injury to the zebrafish retina stimulates MG at the injury site to mount a regenerative response (Fausett & Goldman 2006). This response includes changes in MG gene expression and signal transduction that precedes a self-renewing asymmetric cell division that produces a transient amplifying progenitor for retinal repair (Hoang et al 2020, Lee et al 2020, Nagashima et al 2013, Powell et al 2013, Ramachandran et al 2010a, Ramachandran et al 2011, Ramachandran et al 2012, Sifuentes et al 2016, Wan & Goldman 2017, Wan et al 2012, Wan et al 2014, Zhao et al 2014b).

It remains unknown why mammalian MG cannot mount a regenerative response like their zebrafish counterparts. However, their basal state likely dictates their regenerative/stem cell potential. Recent studies comparing MG transcriptomes across species, as well as targeted perturbation of specific genes and signaling pathways, have identified differences between fish and mammalian MG gene expression programs (Elsaeidi et al 2018, Hoang et al 2020, Lee et al 2020, Sifuentes et al 2016). For example, Notch signaling is active in adult zebrafish MG (Elsaeidi et al 2018, Lee et al 2020, Wan & Goldman 2017, Wan et al 2012), but is dramatically reduced in adult mouse MG (Lee et al 2020, Nelson et al 2011, Riesenberg et al 2018). Interestingly, Notch signaling helps maintain MG quiescence in zebrafish (Campbell et al 2021, Conner et al 2014, Elsaeidi et al 2018, Wan & Goldman 2017, Wan et al 2012).

Using the needle poke injury model we have shown that MG at the injury site engage in a regenerative response that is reflected in their cell division (Fausett & Goldman 2006). This restricted MG response suggests injury signals do not travel far from the injury site and that a certain threshold of injury-derived signal must be exceeded to engage MG in a regenerative response (MG’s injury-response threshold). Indeed, MG do not mount a regenerative response when cell death remains low (Iribarne et al 2019, Lessieur et al 2019, Montgomery et al 2010). A clue to how this injury-response threshold is set comes from the observation that the zone of injury-responsive MG is expanded when a needle poke injury is combined with pan retinal Notch inhibition (Lee et al 2020, Wan & Goldman 2017, Wan et al 2012). Thus, normally quiescent MG near an injury site are recruited to a regenerative response when Notch signaling is inhibited. These data indicate that Notch signaling not only controls the proliferation of MG and MG-derived progenitors (Campbell et al 2021, Conner et al 2014, Lee et al 2020, Wan & Goldman 2017, Wan et al 2012), but also controls MG’s injury-response threshold which represents one of the earliest consequences of MG reprogramming. Understanding the mechanism by which Notch signaling controls MG’s injury-response threshold may provide clues for enhancing the regenerative potential of mammalian MG.

Here we report on studies investigating how Notch signaling regulates MG’s injury-response threshold. We found that Notch signaling inhibition enhances chromatin accessibility and the expression of certain regeneration-associated genes in quiescent MG. Of these genes, two Notch target genes, *hey1* and *id2b*, were identified that mediate the effects of Notch signaling on retina regeneration. Interestingly, Hey1 regulates MG’s injury-response threshold, while both Hey1 and Id2b regulate proliferation of MG-derived progenitors. Furthermore, we found that *hey1* and *id2b* are downstream targets of Dll4/Dlb-Notch3 signaling. Finally, our studies indicate that retention of sufficient Notch signaling that allows for transient suppression distinguishes pro-regenerative MG in zebrafish from non-regenerative MG in mice.

## Results

Although a relatively thorough characterization of MG gene expression and chromatin accessibility accompanying light- or NMDA-induced retinal degeneration was recently reported (Hoang et al 2020), the impact of Notch signaling on these events was not investigated. Importantly, the mechanism by which Notch signaling regulates MG’s responsiveness to injury-derived factors (MG’s injury-response threshold) remains unexplored. We hypothesized that Notch signaling controls MG’s injury-response threshold by impacting chromatin accessibility and gene expression at a subset of regeneration-associated genes. To identify these genes, we searched RNAseq data sets for genes responding to both injury and Notch manipulation, reasoning that this will provide candidate genes that regulate MG’s injury-response threshold and proliferation.

### Injury-dependent regulation of MG gene expression and chromatin accessibility

To identify injury-responsive genes following needle poke injury, we compared RNAseq data sets from GFP+ MG that were FACS purified from the uninjured retina of *gfap:GFP* fish with those from *1016 tuba1a:GFP* fish at 2 days post injury (dpi, 10 needle pokes/retina) (Lee et al 2020). At 2 dpi, most MG are in an activated state with some beginning to undergo cell division (Fausett & Goldman 2006). Differential gene expression analysis identified 4,425 injury-responsive genes with 2,579 showing increased expression (fold change ≥ ± 1.5; Figure 1A). A small subset of these genes were validated by PCR (Figures 1B and S1A). Some of these genes, like *ascl1a* and *hbegfa*, were previously shown to regulate injury-dependent MG proliferation and retina regeneration (Fausett et al 2008, Gorsuch et al 2017, Ramachandran et al 2010a, Ramachandran et al 2011, Wan et al 2012).

**Figure 1:**
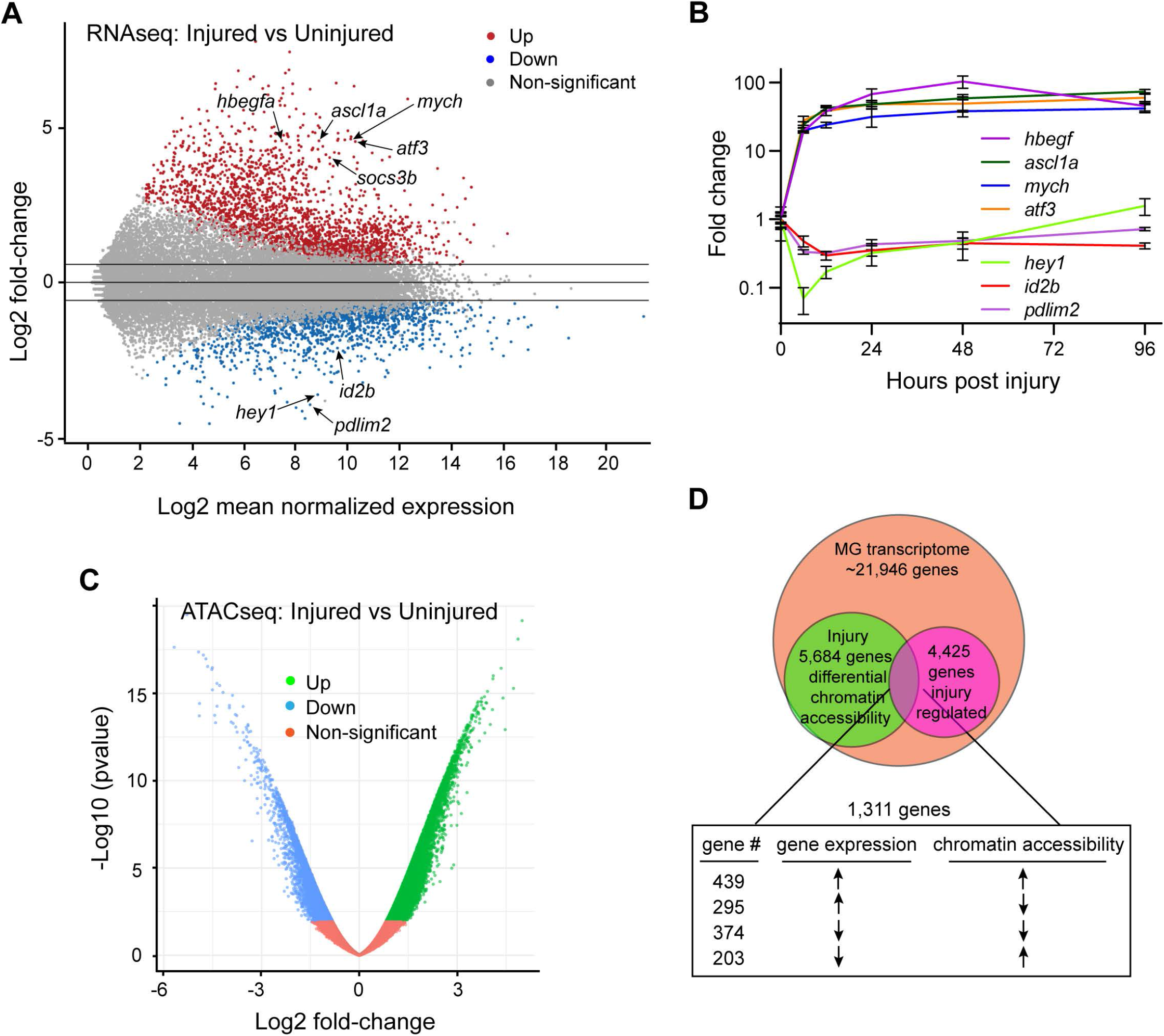
Injury-dependent regulation of MG gene expression and chromatin accessibility. **A**, MA plot of RNAseq data using MG purified from uninjured and injured fish retinas. **B**, qPCR analysis of select injury-regulated genes. **C**, Volcano plot of ATACseq data using MG purified from uninjured and injured fish retinas. **D**, Intersection of regeneration-associated genes with those whose chromatin accessibility changes with injury.

ATACseq was used to assay chromatin accessibility changes as MG transition from a quiescent to an activated state following retinal injury. For this, GFP+ MG and GFP+ activated MG were FACS purified from uninjured *gfap:GFP* fish and injured *1016 tuba1a:GFP* fish (2 dpi, 10 needle pokes/retina), respectively, and nuclei were used for ATACseq (Buenrostro et al 2015). We mapped injury-dependent differential chromatin accessibility to 5,684 genes (Figure 1C). On a global scale, ATACseq revealed a slight bias towards open chromatin in activated MG with 15,595 regions exhibiting an increase and 13,049 regions exhibiting a decrease in chromatin accessibility. Quantification of the number of unique annotations for each region and annotating these regions to different genic features revealed increased chromatin accessibility at intergenic regions and decreased chromatin accessibility at 5’ UTR, promoters, and 1-5 kb region upstream of transcription start sites (Figure S1B).

To correlate injury-dependent changes in gene expression with chromatin accessibility, we intersected our RNAseq and ATACseq injury-regulated gene lists. This analysis identified 1,311 injury-responsive MG genes with changes in chromatin accessibility (Figures 1D and S1C). ~60% of the injury-induced genes and ~64% of the injury-repressed genes showed a corresponding change in chromatin accessibility (Figure 1D; Table S1). Importantly, many of these changes in chromatin accessibility included promoters and upstream sequences (Figures 1 and S1C; Table S2).

### DAPT-dependent regulation of MG gene expression and chromatin accessibility

Notch repression with DAPT allows normally quiescent MG adjacent to the injury affected zone to mount a regenerative response(Wan & Goldman 2017, Wan et al 2012). To understand how Notch inhibition accomplishes this, we used RNAseq to characterize the DAPT-regulated transcriptome in quiescent MG. For this, GFP+ MG were FACS purified from uninjured *gfap:GFP* fish treated +/-DAPT for 24 hours. RNAseq identified 1,261 differentially expressed genes, of which 375 were reduced following Notch inhibition and 245 were also regulated by retinal injury (fold change ≥ ± 1.5; Figure 2A). Interestingly, these latter genes include *ascl1a* and *hbegfa* which are known to be essential for retina regeneration (Figures 2A, 2B, S2A and S2B; Table S3) (Ramachandran et al 2010a, Ramachandran et al 2011, Wan et al 2012).

**Figure 2:**
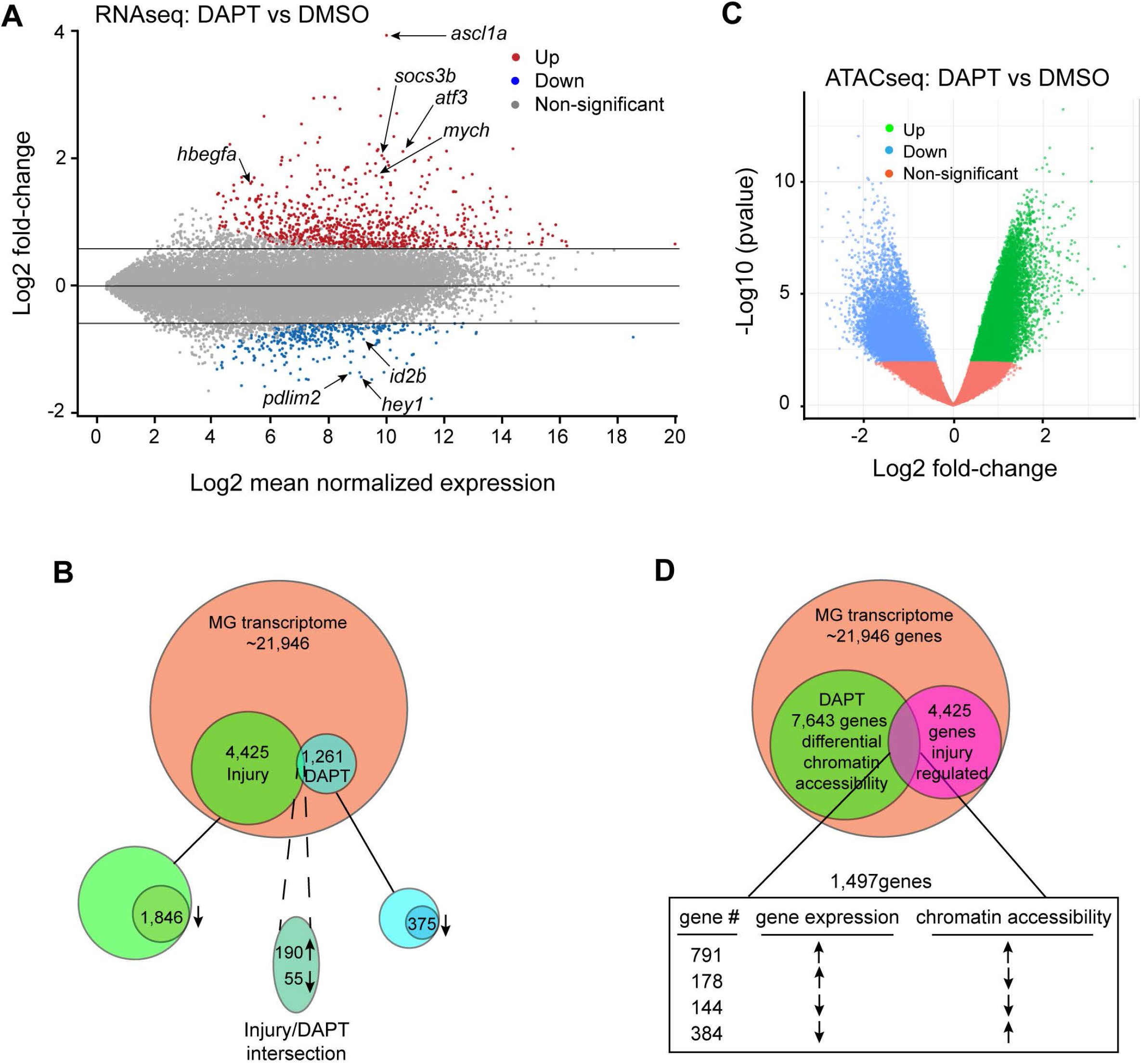
Notch-dependent regulation of MG gene expression and chromatin accessibility in uninjured retina. **A**, MA plot of RNAseq data from MG of uninjured fish treated +/- DAPT. **B**, Intersection of regeneration-associated genes with DAPT-regulated genes. **C**, Volcano plot of ATACseq data from MG of uninjured fish treated +/- DAPT. **D**, Intersection of regeneration-associated genes with those whose chromatin accessibility changes with DAPT treatment.

We next used ATACseq to investigated the consequence of Notch signaling inhibition on MG chromatin accessibility (Buenrostro et al 2015). For this, nuclei were isolated from GFP+ MG purified from uninjured *gfap:GFP* fish treated +/-DAPT. On a global scale, inhibition of Notch signaling affected chromatin accessibility in a similar fashion as retinal injury with 16,990 regions showing increased and 12,413 regions showing decreased chromatin accessibility (Figure 2C). However, unlike that found with retinal injury, quantification of the number of unique annotations for each region and annotating these regions to different genic features revealed that DAPT-treatment increased chromatin accessibility in 5’ regions of genes including 5’UTR, promoters, and 1-5kb upstream regions, while intergenic regions appear to be enriched for closed chromatin (Figure S2C and S2D).

We mapped DAPT-dependent differential chromatin accessibility to 7,643 genes (Figure 2D), which exceeds those whose chromatin accessibility is regulated by retinal injury (Figure 1D). Interestingly, when examining promoter regions (1kb upstream of transcription start site) for differential chromatin accessibility, we found 156 promoters with reduced chromatin accessibility, and 3,871 promoters with increased chromatin accessibility (Table S4). This bias towards open chromatin was also observed in the 1-5kb region upstream of transcription start sites, where we observed 196 genes with reduced chromatin accessibility and 1,041 genes with increased chromatin accessibility (Table S4).

Of the genes whose chromatin accessibility changed with DAPT-treatment, the expression of 1,497 were also regulated by retinal injury (Figure 2D), which is similar to the number of genes showing changes in chromatin accessibility after retinal injury (Figure 1D). However, when we correlate injury-dependent gene regulation with changes in chromatin accessibility upstream of transcription start sites, we found a larger proportion of the injury-responsive genomic regions with increased chromatin accessibility following DAPT-treatment compared to retinal injury (1,175 vs 642) (Figures 1D and 2D; Tables S2 and S4). Furthermore, ~67% of these latter increases in chromatin accessibility are associated with genes whose expression is increased in the injured retina.

Together, the above data suggests Notch inhibition in the uninjured retina contributes to lowering MG’s injury-response threshold by increasing chromatin accessibility and regulating the expression of certain key regeneration-associated genes.

### Notch-dependent regulation of genes impacting MG reprogramming

We hypothesized that MG regeneration-associated genes that are also regulated in a similar direction by Notch signaling in the uninjured retina would identify a relatively small set of genes that may regulate MG’s injury-response threshold. Intersection of injury and Notch-regulated RNAseq data sets revealed 190 induced (basal value ≥5; fold change ≥1.5) and 55 repressed (basal value ≥50; fold change ≥1.5) genes (Figures 2B, S2A, and S2B; Table S5). The top 20 injury-responsive genes that were regulated in similar direction following DAPT-treatment are listed in Figure S2A.

DAPT is a γ-secretase inhibitor that can target over 100 substrates, of which Notch is only one (Xia 2019). To help narrow-in on Notch-regulated genes, we took advantage of *hsp70:DN-MAML* transgenic fish (Zhao et al 2014a). DN-MAML suppresses endogenous Notch-dependent gene activation (Maillard et al 2006, Maillard et al 2004). To identify MG genes regulated by DN-MAML in uninjured retina, we treated *gfap:mCherry* and *gfap:mCherry;hsp70:DN-MAML* fish with heat shock every 6 hours for 1 day and then FACS purified GFP+ MG for RNAseq. Expression analysis identified 1,122 differentially expressed genes with only 244 showing reduced expression (fold change ≥ ± 1.5; Figure S2E).

We next determined the intersection of injury-responsive MG genes that were also regulated in the same direction by DAPT and DN-MAML in the uninjured retina. Remarkably, this analysis identified 84 induced genes and 5 repressed genes (fold change ≥ ± 1.5) (Table S5 and genes marked with asterisk in Figure S2A). Thus, out of a total of 4,425 regeneration-associated genes, we have narrowed-in on 89 that are Notch-responsive in the uninjured retina and perhaps regulating MG’s injury-response threshold.

### Hey1 and Id2b regulate MG’s injury-response threshold and proliferation

*hey1* and *id2b* were the only transcriptional regulators consistently repressed with retinal injury or Notch inhibition, making them good candidates for being direct Notch target genes. Inspection of the 3kb region upstream of their transcriptional start sites revealed consensus Rbpj binding elements in the *hey1* gene (Figure 3A). Two of these elements are highly conserved in sequence and position when compared with orthologous genes from mouse and human (Figure 3A). Mutation of either of these putative Rbpj binding sites in a 3kb *hey1* promoter fragment driving luciferase gene expression resulted in a dramatic reduction in promoter activitation by NICD when assayed in transfected HEK293 cells (Figure 3A and 3B).

**Figure 3:**
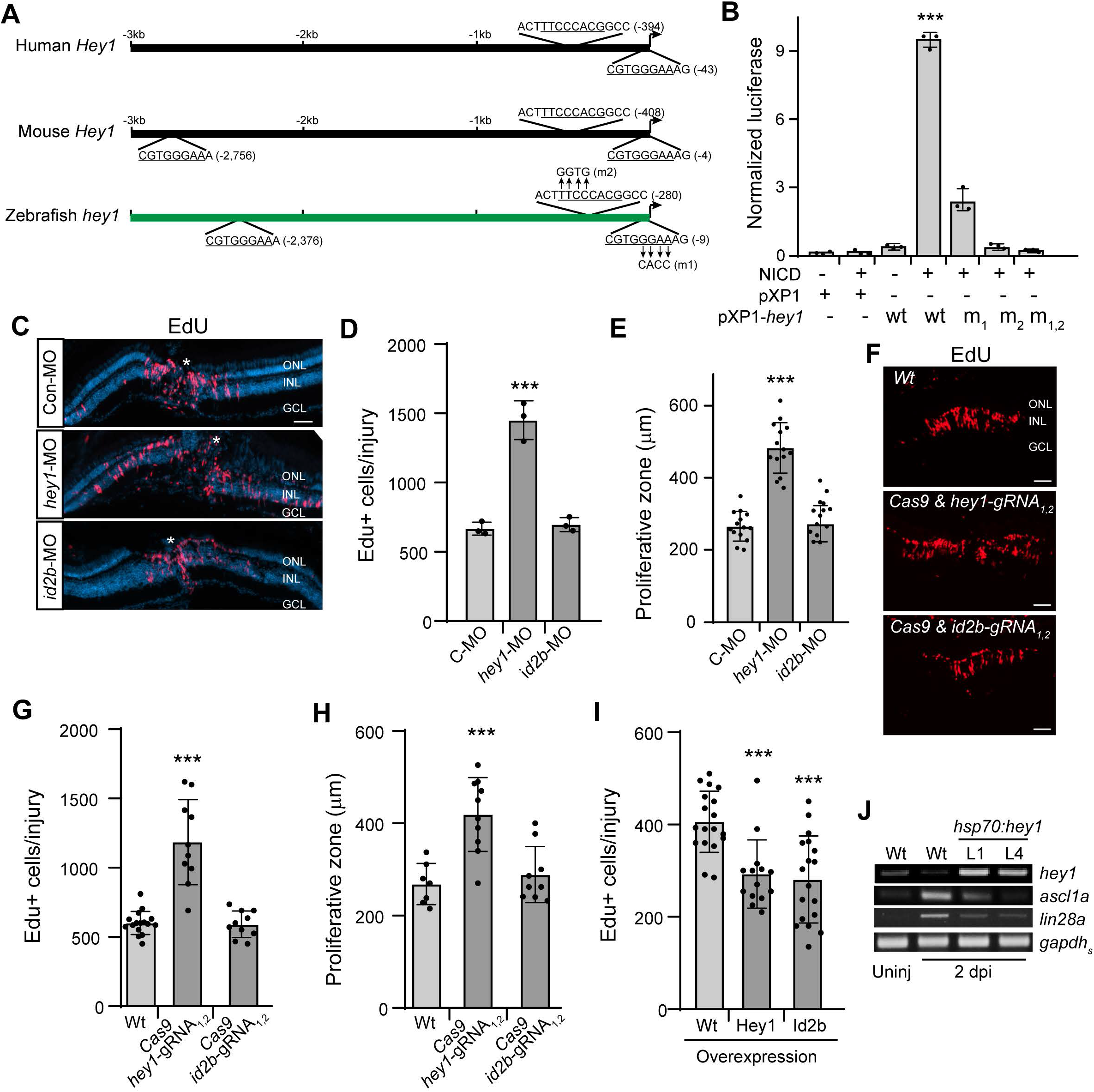
Hey1 and Id2b regulate MG’s regenerative response. **A**, Illustration of *hey1* gene promoter from human, mouse and zebrafish with Rbpj binding sites indicated. **B**, Luciferase assays for Notch (NICD) dependent activation of Wt and mutant *hey1* promoter activity; Compared to Wt alone, \*\*\**P*<0.0001. **C**, **F**, EdU Click-iT chemistry on retinal sections identifies proliferating cells in Wt fish treated with indicated MOs (**C**) or Cas9 and indicated gRNAs (**F**). **D**, **G**, **I**, Quantification of the number of EdU+ cells at 4 dpi; (**D**)\*\*\**P*=0.0008, (**G**) \*\*\**P*<0.0001, (**I**) \*\*\**P*<0.0001. **E**, **H**, Quantification of the width of the proliferative zone in retinas at 4 dpi; (**E**) \*\*\**P*<0.0001, (**H**) \*\*\**P*=0.0004. **J**, PCR analysis of the effects Hey1 expression has on injury-dependent induction of *ascl1a* and *lin28a*. Abbreviations: MO, morpholino; C-MO or Con-MO, control-MO; ONL, outer nuclear layer; INL, inner nuclear layer; GCL, ganglion cell layer; Wt, wild type; m, mutant. Scale bars are 100 μm.

To investigate if Hey1 and Id2b contributed to MG’s injury-response threshold, we knocked down their expression with either a splice blocking *hey1*-morpholino (MO) or a translation blocking *id2b*-MO (Figure S3A-C). MO effectiveness was determined in embryos, where injection of the *hey1*-MO resulted in *hey1* RNA intron-retention and injection of the *id2b*-MO with an *id2b*-GFP chimeric RNA resulted in reduced GFP expression (Figure S3B and S3C). Remarkably, only Hey1 knockdown expanded the zone of injury-responsive MG (Figures 3C-E, and S3D). We confirmed this result using a CRISPR-based gene editing strategy where embryos were injected with *in vitro* transcribed *Cas9-nanos* and 2 different gRNAs targeting either *hey1* or *id2b* exons 1 and 2 (Figure S3A). These fish were raised to adults and gene deletions were confirmed using genomic DNA from fin clips (Figure S3E and F). Consistent with our knockdown experiments, only editing of the *hey1* gene resulted in an expanded zone of injury-responsive MG (Figure 3F-H), which was not associated with changes in retinal cell death (Figure S3G).

Although only Hey1 regulated MG’s injury-response threshold, both Hey1 and Id2b suppressed MG proliferation without affecting cell death in *hsp70:GFP-P2A-hey1* or *hsp70:id2b-P2A-GFP* fish (Figure 3I; Figure S3H). Hey1 overexpression also suppressed injury-dependent induction of regeneration-associated genes like *ascl1a* and *lin28a* (Figure 3J).

Together, the above data show that Notch-regulated genes affecting MG proliferation, like *id2b*, do not necessarily affect MG’s injury-response threshold. However, genes that regulate MG’s injury-response threshold, like *hey1*, also regulate the number of MG engaged in a proliferative response. These data suggest *hey1* and *id2b* reflect a bifurcation in Notch signaling.

### Injury-dependent changes in Notch signaling component gene expression

We previously reported that *hey1* repression correlates with *dll4* repression in the injured retina (Wan & Goldman 2017). Thus, based on the above data, Dll4 may regulate Notch signaling and MG’s injury-response threshold. Indeed, of all Notch ligand encoding RNAs expressed in the retina, *dll4* is the most abundant and its injury-dependent suppression exceeds that of any other Notch ligand encoding gene (Figures 4A 4B, and S4A)(Campbell et al 2021, Hoang et al 2020). Importantly, this suppression temporally follows Notch signaling repression (Figures 4B, and S4E). However, contrary to the idea that Dll4 regulates MG’s injury-response threshold is the observation that Dll4 knockdown inhibited proliferation of MG-derived progenitors in the light damaged retina, while Dlb knockdown enhanced this proliferation (Campbell et al 2021). Further support for Dlb regulating MG quiescence is the finding that *dlb* is enriched in the neuronal population while *notch3* is enriched in the MG population, suggesting Dlb-Notch3 signaling controls MG quiescence and the suppression of this pathway allows for MG proliferation(Campbell et al 2021).

**Figure 4:**
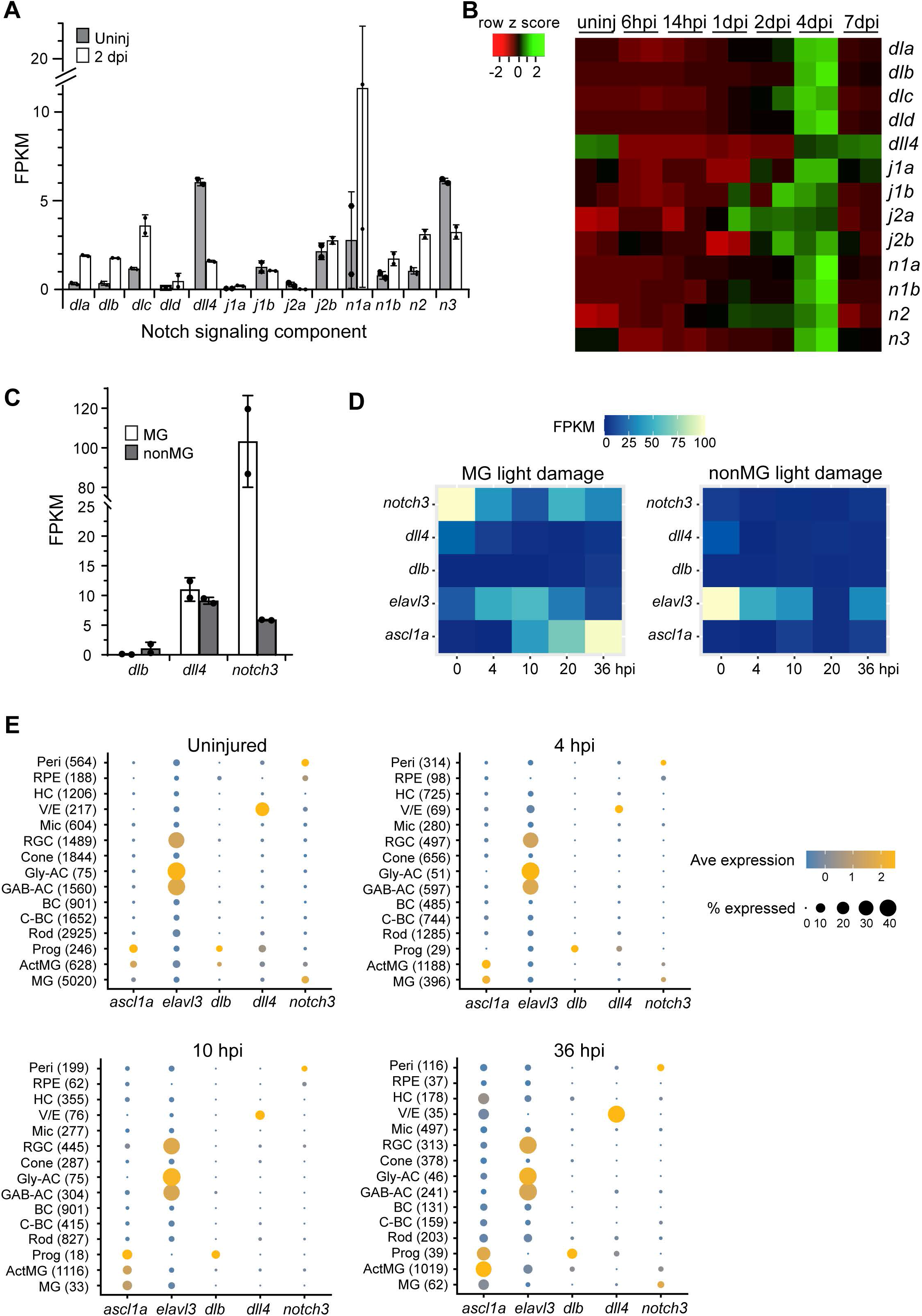
Analysis of Notch ligand and receptor gene expression in the uninjured and injured zebrafish retina. **A**, Expression level of indicated RNAs based on bulk RNAseq data generated from GFP+ MG FACS purified from uninjured *gfap:GFP* and GFP+ activated MG and MG-derived progenitors FACS purified from injured (2 dpi) *1016 tuba1a:GFP* fish(Lee et al 2020). **B**, qPCR heat map illustrating temporal changes in the expression of Notch ligand and receptor encoding genes at various times post needle poke injury. **C**, Expression level of indicated RNAs in MG and nonMG based on bulk RNAseq data generated by Hoang et al., 2020 using GFP+ MG and GFP-nonMG FACS purified from uninjured *gfap:GFP* fish. **D**, Temporal changes in indicated genes at various times post light damage based on bulk RNAseq data generated by Hoang et al., 2020 using GFP+ MG and GFP-nonMG FACS purified from injured *gfap:GFP* fish. **E**, Dot plot visualization of gene expression across retinal cell types in the uninjured and NMDA injured retina based on scRNAseq data from Hoang et al., 2020. Abbreviations: j, jag; n, notch; hpi, hours post injury; dpi, days post injury; FPKM, Fragments Per Kilobase of transcript per Million mapped reads; peri, pericyte; RPE, retinal pigment epithelium; HC, horizontal cell; V/E, vascular endothelial cell; Mic, microglia; RGC, retinal ganglion cell; Gly-AC, glycinergic amacrine cell; GAB-AC, GABAergic amacrine cell; BC, bipolar cell; C-BC, cone bipolar cell; Prog, progenitors in the ciliary margin; ActMG, activated MG.

We were puzzled by the finding that *dlb*, whose overall basal expression and injury-dependent suppression is less than that of *dll4* would be the predominant regulator of Notch signaling in quiescent MG. One possibility is that *dlb* expression exceeds that of dll4 in a pan neuronal fashion, while dll4 expression is restricted to a subset of neurons and/or other cell types. Therefore, we queried previously generated bulk RNAseq data sets from MG and nonMG, and scRNAseq data sets from uninjured and light or NMDA-damaged retina(Hoang et al 2020). This analysis revealed that *dll4* expression exceeds that of *dlb* in both MG and nonMG populations and that *notch3* was enriched in the MG population (Figure 4C). Importantly, retinal injury suppressed *dll4* in both the MG and nonMG population, while *dlb* was less affected (Figure 4D). As controls, we evaluated *elavl3* and *ascl1a* expression - *elavl3* is an amacrine and ganglion cell marker, while *ascl1a* is a marker of activated and proliferating MG and MG-derived progenitors (Figure 4D).

Further interrogation of scRNAseq data sets indicated *dlb* and *dll4* are expressed at low levels in MG, resulting in only a small fraction of MG exhibiting expression above background (Figures 4E and S4B). Nonetheless, in the uninjured retina, *dlb* is enriched in progenitors located in the ciliary margin and activated MG, while *dll4* predominates in the vascular endothelial (V/E) cell population (Figures 4E and S4b). Both genes are poorly expressed in neurons; however, *dll4* expression exceeds that of *dlb* (Figure 4E and S4B). Importantly, retinal injury rapidly reduced *dll4* expression in V/E cells, while *dlb* expression was reduced in activated MG (Figure 4E). In contrast, *notch3* expression is highest in the pericyte and MG population and this MG expression is reduced after injury (Figure 4E). As expected, scRNAseq data sets show induction of *ascl1a* in activated MG following retinal injury, while *elavl3* expression is constitutively expressed in amacrine (AC) and retinal ganglion cell (RGC) populations (Figure 4E). The increased expression of *ascl1a* in resting MG after injury indicates that a fraction of these cells are in an activated state.

The above analysis raises the interesting possibility that neurodegeneration in the retina impacts V/E cells to control *dll4* expression. The relatively high basal expression of *dll4* in V/E cells and its coordinate repression with *notch3* and Notch signaling in MG, suggests a mechanism for regulating Notch signaling in MG. Indeed, MG end-feet are in direct contact with V/E cells (Alvarez et al 2007). Importantly, both photoreceptor degeneration in the light damaged retina and degeneration of amacrine and RGCs in the NMDA damaged retina reduce *dll4* expression in V/E cells (Figure 4E) (Hoang et al 2020). However, cell type-specific *dll4* and *dlb* gene knockdown or inactivation in MG, V/E cells, and neurons is necessary to confidently identify the cell sources controlling Notch signaling in MG.

### Notch signaling inhibits proliferation of MG-derived progenitors

The above studies presented a conundrum in that *dll4* gene expression suggested it might contribute to the maintenance of Notch signaling in quiescent MG, yet its knockdown was reported to inhibit proliferation of MG-derived progenitors (Campbell et al 2021). Furthermore, it seemed odd that knockdown of *dll4* expression, which is already repressed to very low levels after retinal injury, would have a similar effect on proliferation of MG-derived progenitors as repression of *dla, dlc*, or *dld* whose expression is highly induced in the injured retina (Figures 4A, 4B, and S4A) (Campbell et al 2021). Finally, this latter result is at odds with a previous study showing conditional expression of NICD at 3-4 dpi inhibited proliferation of MG-derived progenitors (Wan et al 2012), which is consistent with Notch signaling inhibiting proliferation or MG-derived progenitors. Thus, a further analysis of the role Dll4 and Notch signaling play in retina regeneration seemed warranted.

To further explore the relationship between Notch signaling and proliferation of MG-derived progenitors, we injured fish retinas with needle poke and labelled proliferating cells at 2 dpi with an intraperitoneal (IP) injection of EdU. Fish were then immersed in either DMSO or DAPT for an additional 2 days before receiving an IP injection of BrdU 3hrs prior to sacrifice. Quantification of the fraction of EdU+ cells that continued to proliferate at 4 dpi (BrdU+ & EdU+) revealed enhanced proliferation of MG-derived progenitors when Notch signaling was inhibited (Figure S4F). This result is most consistent with the idea that Notch signaling promotes quiescence of MG and MG-derived progenitors. Thus, we propose that the return of Notch signaling to the injured retina around 4 dpi (Figure S4E), which parallels the large increase in expression of *dla, dlc*, and *dld* (Figures 4B and S4A), facilitates progenitor cell cycle exit and differentiation.

### Dlb, Dll4 and Notch3 regulate MG’s injury-response threshold

Based on the above data, we hypothesized that Dll4 is a major regulator of Notch signaling in quiescent MG and thereby, regulates MG’s injury-response threshold. To investigate this, we knocked down Dll4 using a *dll4*-MO that blocks *dll4* splicing and results in intron retention (*dll4*-MO, Figures 5A and S5H). Indeed, Dll4 knockdown increased MG proliferation and expanded the zone of proliferating MG in a concentration-dependent manner (Figure 5B-D). Dll4 knockdown had no significant effect on retinal cell death (Figure 5E). In light of previous data indicating Dll4 knockdown stimulates proliferation of MG-derived progenitors (Campbell et al 2021), it was important to verify this result with an independent method. For this we used a CRISPR-based strategy to edit the *dll4* gene. gRNAs were designed to target *dll4’s* DSL domain that interacts with the Notch receptor extracellular domain (gRNAs 1 and 2, Figure 5a). gRNAs were cloned into a Tol2 vector that allows expression of multiple gRNAs from different *u6* promoters (Yin et al 2015), and these fish were named *u6:dll4-gRNA_1,2_*. These fish were bred with *ubi:Cas9* transgenic fish to generate *ubi:Cas9;u6:dll4-gRNA_1,2_* fish. Fish harboring *dll4* gene deletions were identified by PCR using genomic DNA (Figure S5I). Like Dll4 knockdown, retinal injury in *ubi:Cas9;u6:dll4-gRNA_1,2_* fish increased the number of MG engaged in a proliferative response, which was reflected by an expanded zone of injury-responsive proliferating MG (Figure 5F–5H). CRISPR-mediated *dll4* gene editing had no significant effect on cell death (Figure 5I). Finally, forced expression of Dll4 with heat shock in *hsp70:dll4-P2A-GFP* transgenic fish inhibited injury-dependent proliferation of MG and MG-derived progenitors without affecting retinal cell death (Figures 5J–5L and S5J). Together, these data suggest Dll4 regulates MG’s injury-response threshold.

**Figure 5:**
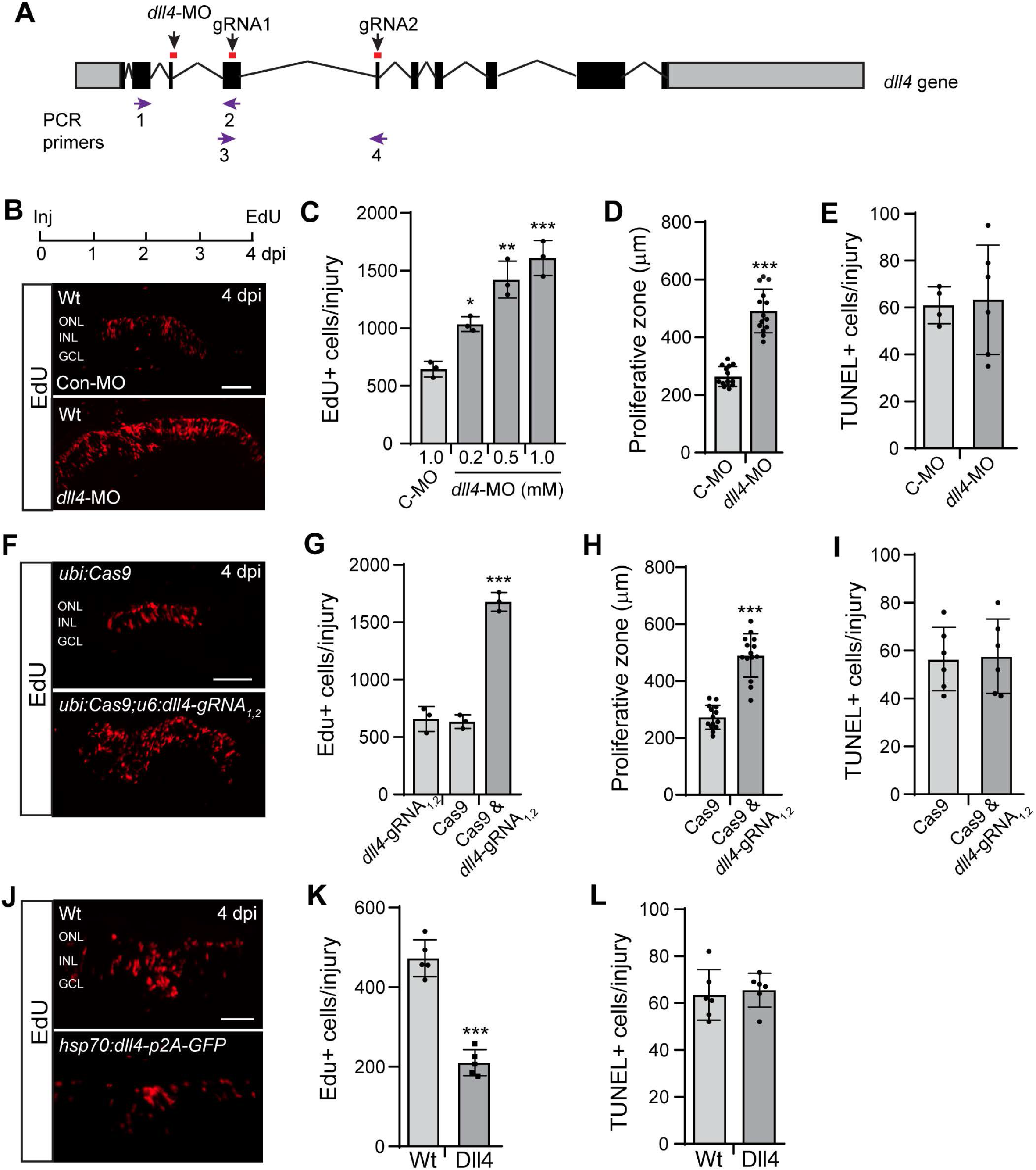
Dll4 regulates MG’s injury response threshold and proliferation of MG-derived progenitors. **A**, Diagram of the *dll4* gene with MO and gRNA target sites indicated. **B, F, J,** Fluorescent images showing EdU Click-iT chemistry on retinal sections at 4 dpi. **C, G, K,** Quantification of the number of EdU+ cells at 4 dpi; (**C**), compared to C-MO\**P*=0.0351 (0.2mM), \*\**P*=0.0022 (0.5mM), \*\*\**P*=0.0003 (1mM); (**G**), compared to Cas9 \*\*\**P*<0.0001; (**K**), \*\*\**P*<0.0001. **D, H,** Quantification of the width of the proliferative zone in retinas at 4 dpi; (**D**), \*\*\**P*<0.0001; (**H**), \*\*\**P*<0.0001. **E, I, L,** Quantification of TUNEL+ cells at 1 dpi. Abbreviations: Wt, wild type; MO, morpholino; C-MO or Con-MO, control-MO; ONL, outer nuclear layer; INL, inner nuclear layer; GCL, ganglion cell layer. Scale bars are 100 μm.

Although basal expression and injury-dependent repression of *dlb* is significantly less than *dll4* (Figure 4), Dlb knockdown results in increased MG proliferation in the injured retina, suggesting it too may contribute to MG’s injury-response threshold (Campbell et al 2021). To directly test this, we used a translation blocking MO to suppress Dlb in the needle poke injured retina. The *dlb*-MO was validated in zebrafish embryos injected with a *dlb*-GFP chimeric transcript (Figure S5A–S5d). Following Dlb knockdown, we noted an expanded zone of proliferating MG (Figure S5B–S5D). Furthermore, Dlb overexpression in *hsp70:GFP-p2A-dlb* fish, inhibited injury-dependent proliferation of MG-derived progenitors (Figure S5F and S5G).

Notch3 knockdown was previously reported to stimulate MG proliferation in the injured retina (Campbell et al 2021), suggesting it may contribute to MG’s injury-response threshold. Indeed, using a translation-blocking MO that suppressed chimeric *notch3-GFP* RNA expression, we observed an expanded zone of injury-responsive MG in the injured retina (Figure S6A–S6F). We confirmed this result using a CRISPR-based strategy to delete the *notch3* gene’s ankyrin repeat domain that is required for recruiting MAML to the Rbpj-NICD complex (gRNA1 and gRNA2, Figure S6A and S6G) (Nam et al 2006). Retinal injury in *ubi:Cas9;u6:n3-gRNA_1,2_* fish resulted in an expanded zone of injury-responsive MG and increased MG proliferation without any changes in retinal cell death (Figure S6H–S6K).

Together, the above data are consistent with the idea that Dll4 and Dlb acting through Notch3 regulate Notch signaling in MG and regulates MG’s injury-response threshold.

### Epistasis experiments suggest bifurcation of Notch signaling via regulated expression of *hey1* and *id2b*

The above studies suggested Dll4 and Dlb engage the Notch3 receptor to stimulate Hey1 and Id2b expression and that Hey1 regulates MG’s injury-response threshold, while Id2b regulates the proliferation of MG-derived progenitors (Figure 3C–3I). To further investigate these gene interactions, we performed epistasis experiments and assayed the zone of injury-responsive MG and MG proliferation at 2 dpi when MG are beginning to proliferate (Fausett & Goldman 2006). Consistent with Notch3 acting downstream of Dll4 and Dlb, we found that Notch3 knockdown expanded the zone of proliferating MG in transgenic fish overexpressing Dll4 or Dlb. This was also reflected by an increase in proliferation of MG and MG-derived progenitors (Figure 6A and 6B). Interestingly, when we tested if Hey1 and Id2b acted downstream of these Notch signaling components, we found only Hey1 was able to suppress the expanded zone of injury-responsive MG resulting from knockdown of Dll4, Dlb, and Notch3, and this was also reflected in reduced proliferation of MG and MG-derived progenitors (Figure 6C and 6D). However, a small effect of Id2b overexpression on EdU+ cells in the control MO treated retina was noted (Figure 6D), which is consistent with Id2b suppressing progenitor proliferation at 4 dpi (Figure 3I).

**Figure 6:**
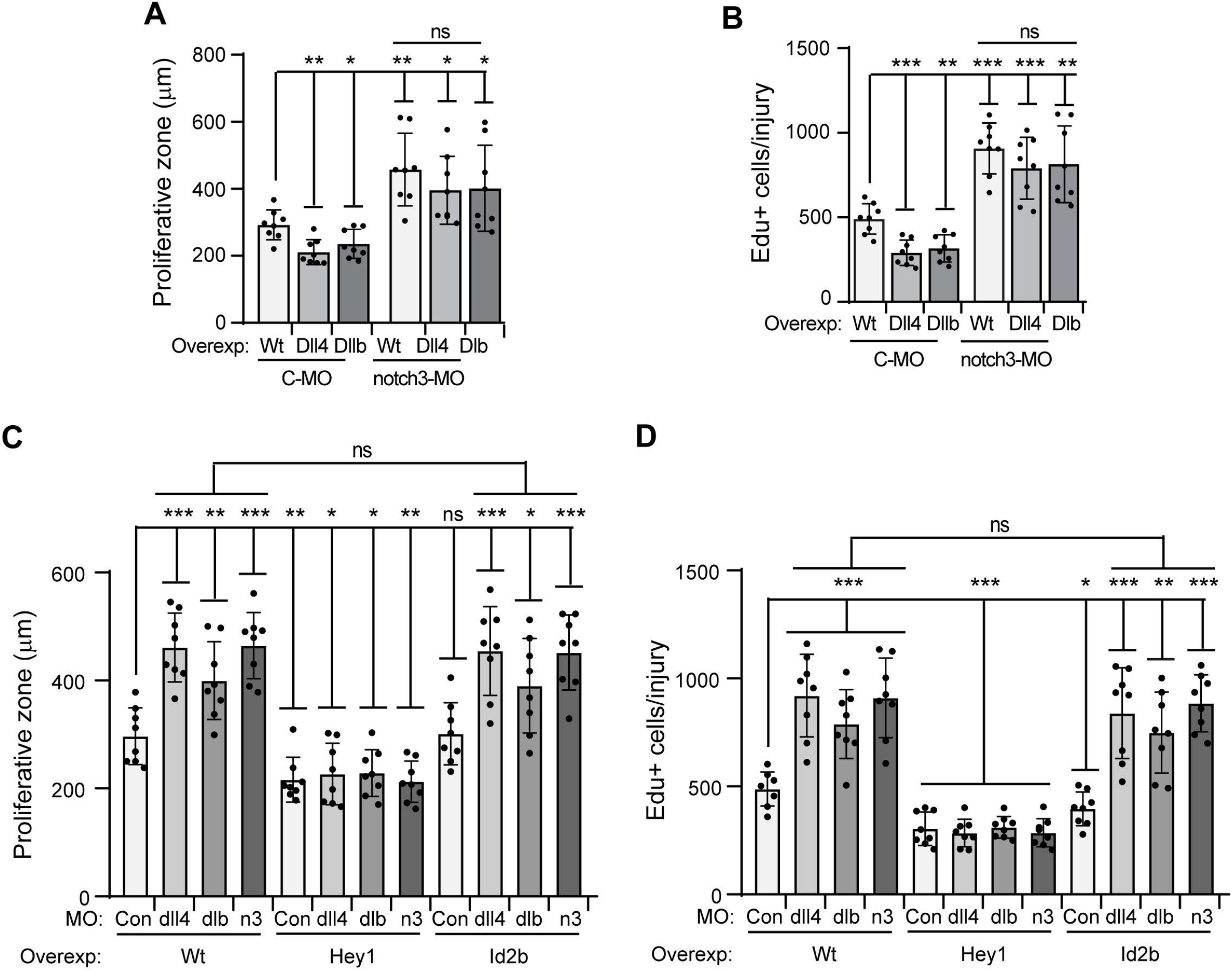
Dll4 and Dlb engage the Notch3 receptor to induce Hey1 and Id2b expression and control retina regeneration. **A, C**, Quantification of the zone of proliferating MG at a needle poke injury site (2 dpi). **B**, **D**, Quantification of proliferating MG-derived progenitors at a needle poke injured site (2 dpi). (**A**), Compared to (C-MO, Wt), \*\**P*=0.0014 (C-MO, Dll4); \**P*=0.0218 (C-MO, Dlb); \*\**P*=0.0031 (notch3-MO, Wt); \**P*=0.0197 (notch3-MO, Dll4); \**P*=0.0394 (notch3-MO, Dlb). (**B**), Compared to (C-MO, Wt), \*\*\**P*=0.0003 C-MO, Dll4); \*\**P*=0.0011 (C-MO, Dlb); \*\*\**P*<0.0001 (notch3-MO, Wt); \*\*\**P*=0.0009 (notch3-MO, Dll4); \*\**P*=0.0022 (notch3-MO, Dlb). (**C**), Compared to control MO (Con) in Wt retina, \*\*\**P*<0.0001 (Wt, dll4-MO); \*\**P*=0.0056 (Wt, dlb-MO); \*\*\**P*<0.0001 (Wt, n3-MO); \*\**P*=0.0042 (Hey1, Con-MO); \**P*=0.0228 (Hey1, dll4-MO); \**P*=0.0132 (Hey1, dlb-MO); \*\**P*=0.0025 (Hey1, n3-MO); \*\*\**P*=0.0004 (Id2b, dll4-MO); \**P*=0.021 (Id2b, dlb-MO); \*\*\**P*=0.0002 (Id2b, n3-MO). (**D**), Compared to control MO (Con) in Wt retina, \*\*\**P*<0.0001 (Wt, dll4-MO); \*\*\**P*=0.0003 (Wt, dlb-MO); \*\*\**P*<0.0001 (Wt, n3-MO); \*\*\**P*=0.0003 (Hey1, Con-MO); \*\*\**P*<0.0001 (Hey1, dll4-MO); \*\*\**P*=0.0001 (Hey1, dlb-MO); \*\*\**P*=0.0001 (Hey1, n3-MO); \**P*=0.0348 (Id2b, Con-MO); \*\*\**P*=0.0006 (Id2b, dl4-MO); \*\**P*=0.0027 (Id2b, dlb-MO); \*\*\**P*<0.0001 (Id2b, n3-MO). Abbreviations: Wt, wild type fish; MO, morpholino; C-MO or con, control MO; n3, notch3; ns, not significant.

Thus, our data are consistent with a Dll4/Dlb-Notch-Hey1 signaling pathway that regulates both MG’s injury response threshold and proliferation, while a Dll4/Dlb-Notch-Id2b signaling pathway appears to predominantly regulate proliferation of MG-derived progenitors. Thus, Hey1 and Id2b reflect a bifurcation of the Notch signaling pathway where downstream effectors have unique and overlapping consequence on retina regeneration.

### Notch signaling in developing and adult zebrafish

In the adult zebrafish retina, Notch signaling is restricted to quiescent MG where it regulates MG’s regenerative response (Elsaeidi et al 2018, Wan & Goldman 2017). In the mammalian retina *Hes5* expression, which is used as a proxy for Notch signaling activity, is readily detected in MG at P7; however, by P30 this expression is reduced to very low levels with endogenous *Hes5* expression reaching the limits of detection (Nelson et al 2011). Furthermore, although *Hes5:GFP* transgenic mice indicate a low level of Hes5 expression in mouse MG at P21-30 (Nelson et al 2011, Riesenberg et al 2018), these very low levels of Hes5 expression are not impacted by inhibitors of Notch signaling (Elsaeidi et al 2018). Importantly, the reduced Notch signaling activity associated with maturing MG in the mouse retina correlates with a loss in their regenerative potential (Loffler et al 2015).

To compare this mammalian pattern of retinal Notch signaling during development with that of fish, we crossed *tp1:mCherry* Notch reporter fish with *gfap:GFP* fish that label MG (Kassen et al 2007, Parsons et al 2009). In developing fish, mCherry and GFP immunofluorescence indicate Notch signaling is activated and restricted to differentiating MG within the first 2 days post-fertilization (dpf), preceding *gfap:GFP* transgene induction (Figure 7a). mCherry immunofluorescence was not detect in progenitors residing in the retinal periphery. PCR analysis of retinal *mCherry* RNA levels indicates a gradual reduction in Notch signaling as the fish matures (Figure 7b and 7c). Nonetheless, unlike that of mouse MG, residual Notch signaling in adult zebrafish MG is readily detected and regulated by retinal injury, γ-secretase inhibitors, and DN-MAML (Figures 7D, and S4C-F,)(Elsaeidi et al 2018, Wan & Goldman 2017). Thus, a major distinction between pro-regenerative MG in zebrafish and non-regenerative MG of mammals is the level of Notch signaling that is retained into adulthood.

**Figure 7:**
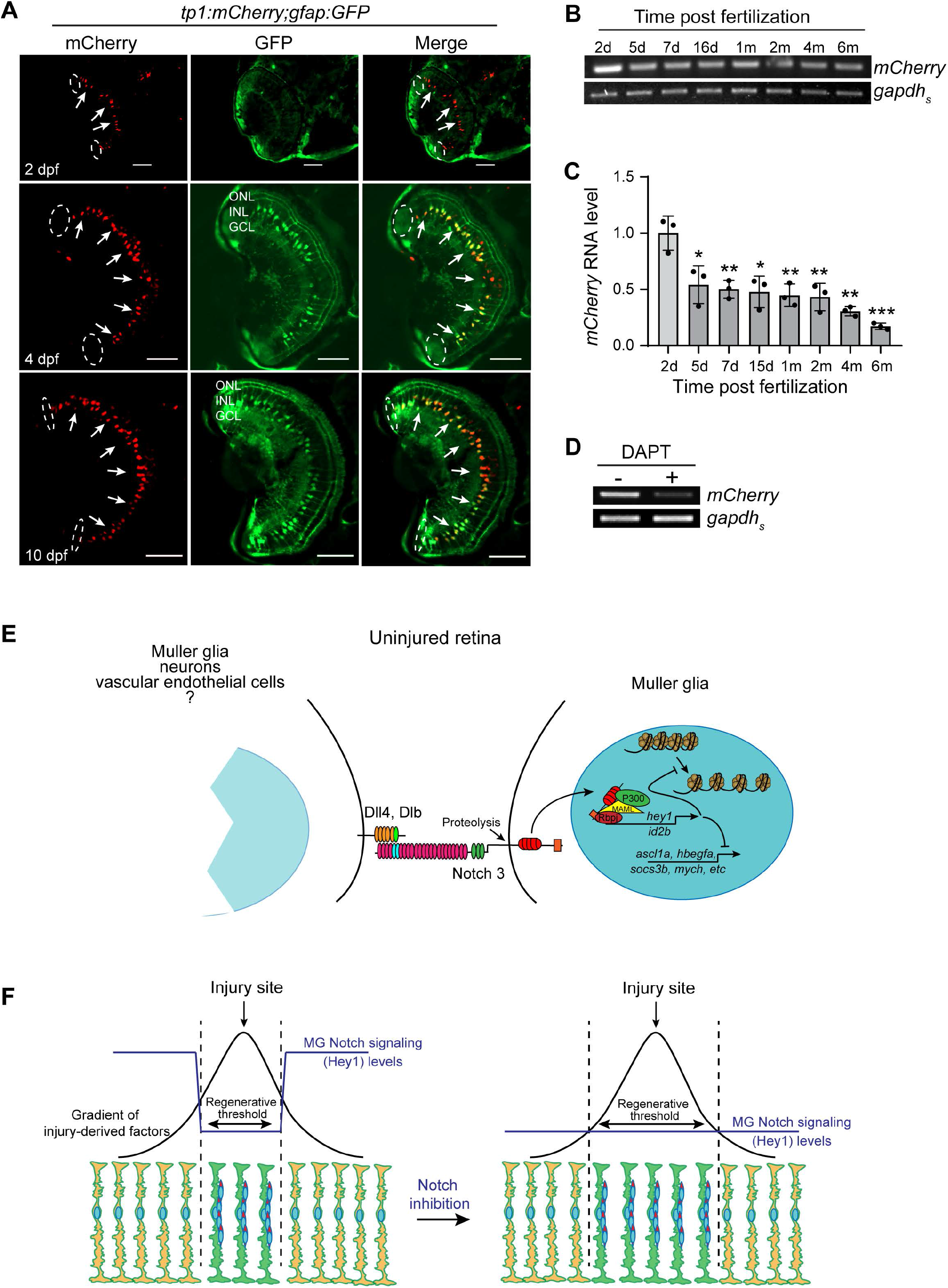
Notch signaling in the developing retina. **A**, GFP and mCherry immunofluorescence on retinal sections from *tp1:mCherry;gfap:GFP* transgenic fish at different times post fertilization. Dashed oval identifies retinal periphery where retinal stem cells involved in retinal expansion reside. **B**, PCR and agarose gel analysis of *mCherry* RNA expression in developing *tp1:mCherry* fish. **C**, qPCR analysis of *mCherry* RNA expression in retinas of developing *tp1:mCherry* fish. Compared to 2 dpf, \**P*=0.0252 (5dpf), \*\**P*=0.0076 (7dpf), \**P*=0.0122 (15dpf), \*\**P*=0.0065 (1mpf), \*\**P*=0.0074 (2mpf), \*\**P*=0.0016 (4mpf), \*\*\**P*=0.0008 (6mpf). Abbreviations: d, days; m, months. Scale bar is 100 μm. **D**, PCR and agarose gel analysis of *mCherry* and *gapdhs* RNA expression in adult *tp1:mCherry* fish treated +/-DAPT. **E** and **F**, Model illustrating the impact Notch signaling has on MG’s regenerative response. **E,** Illustration of the effect of Notch signaling on MG gene expression and chromatin accessibility. Potential retinal cell sources of Dll4 and Dlb are listed. **B**, Illustration of the effect Notch signaling has on MG’s threshold response to injury-derived factors. Abbreviations: ONL, outer nuclear layer; INL, inner nuclear layer; GCL, ganglion cell layer. Scale bar is 50 μm. Abbreviations: ONL, outer nuclear layer; INL, inner nuclear layer; GCL, ganglion cell layer; dpf, day post fertilization.

## Discussion

In the zebrafish retina, Notch signaling is restricted to quiescent MG. Following retinal injury, Notch signaling is suppressed in MG that engage in a regenerative response. Notch signaling regulates injury-dependent proliferation of MG and MG-derived progenitors (Campbell et al 2021, Conner et al 2014, Elsaeidi et al 2018, Lee et al 2020, Wan & Goldman 2017, Wan et al 2012), while our studies suggest Notch signaling regulates MG’s injury-response threshold (Wan & Goldman 2017, Wan et al 2012). This predicts a gradient of injury-derived factors that must exceed a certain level in order to engage MG in a regenerative response (Fig. 7F). Indeed, low levels of neuron death do not stimulate MG proliferation (Iribarne et al 2019, Lessieur et al 2019, Montgomery et al 2010). Our studies indicate that the amount of injury-related factors needed to elicit a regenerative response is determined in part by the basal level of Notch signaling in MG (Figure 7F).

In this report, we focused on mechanisms underlying these Notch-regulated processes. Our data suggests that Notch signaling reduces chromatin accessibility and expression of a subset of regeneration-associated genes to ensure that MG do not inappropriately enter a regenerative response (Figure 7E). By intersecting MG RNAseq gene lists from injured retina with those from uninjured retina +/- Notch inhibition, we identified Hey1 and Id2b as regeneration-associated, Notch-regulated genes. Our data suggest that Hey1 and Id2b reflect a split in the Notch signaling pathway where Hey1 impacts MG’s injury-response threshold and Id2b impacts proliferation of MG-derived progenitors. In an effort to identify genes that stimulate Notch signaling in the uninjured retina, we focused on the most highly expressed Notch ligand and receptor encoding genes in quiescent MG. This analysis revealed that in addition to Dlb (Campbell et al 2021), Dll4 is a potent regulator of MG’s injury response threshold and proliferation (Figure 7E). We confirmed a Dll4/Dlb-Notch3-Hey1/Id2b signaling cascade using epistasis experiments and found that Notch inhibition from 2-4 dpi enhanced proliferation of MG-derived progenitors. Finally, our data suggest that the level of Notch signaling in MG appears to distinguish pro-regenerative MG of the zebrafish retina from non-regenerative MG of mammals.

We previously reported that Notch inhibition is insufficient to drive MG proliferation in the uninjured retina (Elsaeidi et al 2018), and our analysis of Notch-regulated gene expression in quiescent MG suggests this may result from incomplete activation of the regeneration-associated transcriptome. However, when combined with other factors like Tnfa or Ascl1a and Lin28a, MG proliferation is observed (Conner et al 2014, Elsaeidi et al 2018). Thus, Notch signaling repression appears to license MG to engage in a proliferative response, and when exposed to sufficient levels of injury-related factors, MG enter a proliferative phase.

Although Dll4 and Dlb stimulate Notch signaling in quiescent MG, we suspect Dll4 may be the main endogenous factor regulating this process due to its relatively high basal expression, its robust repression following injury, and its better correlation with Notch signaling levels. To identify the cell types expressing these ligands, we queried bulk and scRNAseq data sets and found that both genes are detected at low levels in MG, activated MG, and neurons. However, we show that *dll4* expression generally exceeds that of *dlb* in MG and neurons. Importantly, *dlb* expression predominates in activated MG and progenitor populations, while *dll4* predominates in the V/E population. In addition, injury-dependent suppression of *dll4* was observed in the MG, neuronal, and V/E cell populations, while *dlb* repression was restricted to the activated MG population.

Injury-dependent regulation of *dll4* expression in V/E cells raises the interesting possibility that neuronal injury is communicated indirectly to MG via endothelial cells that make contact with MG end-feet. Indeed, rod loss has been associated with remodeling of the neurovascular unit, as are changes in neural activity (Attwell et al 2010, Garhofer et al 2020, Ivanova et al 2019). However, without cell type-specific gene inactivation we cannot distinguish if Notch signaling in MG is regulated by Notch ligands expressed in neurons, V/E cells, and/or MG.

Our studies indicate *hey1* and *id2b* are downstream targets of Notch signaling in MG. Hey1 is a member of the basic helix-loop-helix orange family of transcriptional repressors, while Id2b is a helix-loop-helix protein that lacks a DNA binding domain and sequesters basic helix-loop-helix proteins, like Ascl1a, from their target genes. Our finding that Hey1’s action downstream of Notch signaling to regulate proliferation of MG and MG-derived progenitors is similar to what’s observed in brain radial glia where Notch signaling maintains neural stem cell quiescence and regulates proliferation of transient amplifying cells (Alunni et al 2013, Engler et al 2018, Imayoshi et al 2010, Kawai et al 2017, Than-Trong et al 2018). Furthermore, Hey1 can suppress expression of Ascl1 and Ascl1-regulated genes which is necessary for establishing a transient amplifying population of neural progenitors in the adult brain (Kim et al 2007, Sakamoto et al 2003). Importantly, Ascl1a is a pioneer transcription factor that can enhance chromatin accessibility across the genome and thereby influencing the expression of many other genes (Wapinski et al 2017). Thus, the Notch-Hey1-Ascl1a signaling axis identified in zebrafish MG appears to be a conserved pathway in a variety of adult radial glial stem cell populations.

In both zebrafish and mice, Notch signaling is necessary for MG differentiation (Bernardos et al 2005, Jadhav et al 2006, Nelson et al 2011, Scheer et al 2001). Notch signaling is restricted to MG in the postnatal mouse retina, and here we report it is also restricted to MG in the developing zebrafish retina. However, while the adult zebrafish retina retains relatively high levels of Notch signaling that is regulated by Notch signaling inhibitors or retinal injury, Notch signaling is suppressed in the adult mouse retina and exhibits no regulation by Notch inhibitors or retinal injury (Elsaeidi et al 2018, Nelson et al 2011). Interestingly, the suppression of Notch signaling during mouse development correlates with the loss of MG stem cell potential (Loffler et al 2015).

In addition to influencing MG’s injury-response threshold, our data supports the idea that Notch signaling inhibition helps maintain MG-derived progenitors in a proliferative state and that the return of Notch signaling around 4 dpi participates in cell cycle exit as progenitors prepare for differentiation. Consistent with this, we previously reported conditional activation of Notch signaling from 3-4 dpi using *hsp70:gal4;uas:nicd-myc* transgenic fish suppressed proliferation of MG-derived progenitors (Wan et al 2012), and here we report that Notch inhibition from 2-4 dpi with DAPT stimulated proliferation of MG-derived progenitors. Furthermore, forced expression of Dll4, Dlb, Notch3, Hey1 or Id2b inhibited proliferation of MG-derived progenitors, while inactivation of these genes enhanced this proliferation. Although these results are all internally consistent with the idea that Notch signaling inhibits proliferation of MG-derived progenitors, Campbell et al., 2021 has come to the opposite conclusion solely based on the effect of MO-mediated gene knockdown experiments. We do not know why our results differ from Campbell et al., 2021, except to note that relying on one method of analysis may not reveal confounding effects.

In addition to their role in retina regeneration, Dll4 and Notch3 are required for angiogenesis and oligodendrocyte development, and loss of either gene is lethal (Leslie et al 2007, Zaucker et al 2013). Thus, one may wonder why the *ubi:Cas9;u6:dll4-gRNAs_1,2_* and *ubi:Cas9;u6:n3-gRNAs_1,2_* transgenic fish used in our studies survive to adulthood. We suspect mosaicism in CRISPR-mediated gene editing underlies this survival. This mosaicism may allow normal development to ensue; however, gene edits will continue to accumulate throughout the fish’s life, which led to the noted enhanced regeneration phenotype in adult fish. We also note that although many homozygous *notch3* mutant fish do not survive to adulthood, some do (Zaucker et al 2013).

In summary, our studies reveal a role for Notch signaling in regulating MG’s injury-response threshold, licensing MG for proliferation, and regulating the proliferation of MG-derived progenitors. Our analysis of mechanisms underlying these processes identified Hey1 and Id2b that reflect a bifurcation point in MG Notch signaling and distinguishes MG’s injury-response threshold from proliferation of MG-derived progenitors. These studies not only further our understanding of how Notch signaling controls retina regeneration in zebrafish, but also suggests that enhancing Notch signaling in mammalian MG may improve their regenerative potential.

## Materials and methods

### Experimental models

Zebrafish: Animal studies were approved by the University of Michigan’s Institutional Animal Care and Use Committee. Zebrafish were kept at 26-28°C with a 14/10 hr light/dark cycle. Adult male and female fish from 6 to 12 months of age were used in these studies. *1016 tuba1a:GFP, gfap:GFP, hsp70:DN-MAML-GFP, gfap:mCherry* and *tp1:mCherry*, fish were previously described(Fausett & Goldman 2006, Johnson et al 2016, Kassen et al 2007, Parsons et al 2009, Zhao et al 2014a). We generated *hsp70:dll4-P2A-GFP, hsp70:GFP-P2A-dlb, hsp70:GFP-P2A-hey1, hsp70:id2b-P2A-GFP, ubi:Cas9, u6:dll4-gRNAs_1,2_* and *u6:n3-gRNAs_1,2_* transgenic fish using standard recombinant DNA techniques and Tol2 vector backbone. Expression constructs were injected into single cell zebrafish embryos as previously described(Fausett & Goldman 2006). Fish were anesthetized in tricaine and retinas were injured with needle pokes (2-4 injuries/retina for analysis of proliferation and protein expression on retinal sections and 8 injuries/retina for qPCR, ATACseq and RNAseq), as previously described(Fausett & Goldman 2006).

### Cell proliferation assays

To investigate cell proliferation, fish received an IP injection of EdU (10 μl of 10mg/ml stock) and Click-It chemistry was used for detection as previously described(Wan & Goldman 2017). Some fish received an IP injection of BrdU (20 mM).

### RNA isolation and PCR

Total RNA was isolated using Trizol (Invitrogen). cDNA synthesis and PCR reactions were performed as previously described(Fausett et al 2008, Ramachandran et al 2010a). Real-time qPCR reactions were carried out using Green 2x LO-ROX qPCR mix (ASI) on an iCycler real-time PCR detection system (BioRad). The ΔΔCt method was used to determine relative expression of mRNAs in control and injured retinas and normalized to either *gapdh* or *gapdhs* mRNA levels. Individual comparisons were done using unpaired 2-tailed Student t-test. ANOVA with Fisher’s PLSD post hoc analysis was used for multiple parameter comparison. Error bars are standard deviation (SD).

### ATACseq and RNAseq

For ATACseq and RNAseq, retinas from injured (2 dpi) *1016 tuba1a:GFP* and uninjured *gfap:GFP* fish were dissociated and GFP+ MG were purified using FACS in the University of Michigan’s Cell Sorting Core as previously described(Powell et al 2013, Ramachandran et al 2010a). Retinas were injured with 10 needle poke injuries and 12 injured *1016 tuba1a:GFP* retinas yielded ~70,000 GFP+ cells. Two retinas from uninjured *gfap:GFP* fish yielded ~225,000 GFP+ cells. For ATACseq nuclei were isolated from GFP+ cells; transposase treatment and PCR amplification were as previously described(Buenrostro et al 2015). For RNAseq, RNA was purified from GFP+ cells and RNA quality and quantity analyzed on a Bioanalyzer (Agilent). cDNA libraries were generated by the U of M’s Advance Genomics Core and DNA was sequenced on an Illumina NovaSeq 6000 platform. Sequencing reads were analyzed by the University of Michigan’s Bioinformatics Core. The number of reads for each expressed gene was determined and differentially expressed genes were restricted to those exhibiting at least a 1.5-fold difference in expression with threshold abundance greater than 5 Fragments Per Kilobase of transcript per Million mapped reads to eliminate very low abundant transcripts whose estimates of fold-change are unreliable.

### Bulk RNA-seq data analysis of MG and nonMG

MG RNAseq data generated from *gfap:GFP* transgenic fish treated +/- NMDA, and light damaged fish retinas were obtained from Hoang et al., 2020. The original raw sequencing reads were processed by the authors as described(Hoang et al 2020). Briefly, RNAseq libraries were prepared from FACS purified MG and nonMG with a result of 45 to 55 million reads per library. Each condition contained two biological replicates, and FPKM (fragments per kilobase of exon per million) were calculated using the raw counts of genes. Using this dataset, we identified genes of interest and the corresponding FPKM across samples were averaged and formulated into heat maps using R for visualization.

### scRNAseq data analysis of control and NMDA-treated retinas

Retinal scRNAseq data from control and NMDA damaged fish retinas were obtained from Hoang et al., 2020. Briefly, the authors used Cell Ranger from 10x Genomics to map zebrafish reads from scRNAseq to the genome with an average of 1000 to 1500 genes and 2500 to 3250 unique molecular identifiers per cell. Variable genes were identified and clustered with K-nearest neighbors and shared nearest neighbor modularity optimization through Seurat. Doublets were eliminated with an in-house algorithm, and marker genes were used to annotate cell types for each cell cluster identified. Using this dataset, we identified the scaled and normalized average expression of our genes of interest across cell clusters, and the percentage of cells per cluster expressing these genes were calculated using R. Dot plots were created using Seurat for visualization.

### CRISPR-based gene editing

Gene editing was performed as previously described(Hwang et al 2013, Vejnar et al 2016). Briefly, we used CRISPRscan to identify 3-6 gRNAs that target the gene of interest(Moreno-Mateos et al 2015). gRNA DNA template synthesis using gRNA primer, universal primer, and *in vitro* transcription are as previously described(Vejnar et al 2016). gRNAs are tested for editing efficiency by injecting single cell embryos with *in vitro* transcribed gRNA, along with Cas9-nanos RNA; 2-4 days later, editing efficiency is determined using genomic DNA in T7 endonuclease 1 (T7E1) mismatch detection assays and PCR(Sentmanat et al 2018). For stable gene editing in transgenic fish, we generated *Ubi:Cas9;cmlc2:GFP* (*Ubi:Cas9*) transgenic fish using standard recombinant DNA protocols and Tol2 vectors. We also generated Tol2 vectors harboring two gRNAs targeting a single gene under the control of two *u6a* promoters as previously described(Yin et al 2015). We chose to express 2 gRNAs for each gene to improve editing efficiency and generate large deletions. Stable transgenic lines were established as previously described and are referred to as *u6:gene-gRNA_1,2_*(Fausett & Goldman 2006). For gene editing in transgenic lines, *Ubi:Cas9* fish and *u6:gene-gRNA_1,2_* were bred to each other and *Ubi:Cas9;u6:gene-gRNA_1,2_* fish (F1 generation) were used for gene editing experiments. Gene editing in transgenic fish was confirmed by PCR and DNA sequencing. For gene editing in F0 fish, we injected zebrafish embryos with previously validated *in vitro* transcribed gRNAs_1,2_ and *Cas9-nanos* mRNA. Embryos were raised to adults and genomic DNA from fin clips was used in T7E1 mismatch detection assay and PCR to confirm gene editing(Sentmanat et al 2018).

### MOs, gRNAs, and qPCR Primers used in this study

All MOs, gRNAs, and primers used in this study are listed in Table Supplement 7.

### Morpholino (MO) functional assays

MOs were obtained from Gene Tools, LLC and the sequences are listed in Supplementary Table 7. All MOs have a lissamine tag. Unless otherwise stated, MOs were used at 1 mM concentration. For testing splice blocking *dll4*-and *hey1*-targeting MO, we injected either control MO or experimental MO (~1ng) into single cell zebrafish embryos. RNA was extracted from embryos at 24-48hrs post injection and assayed for *dll4* or *hey1* mRNA by PCR. For testing *dlb*-, *n3*-, and *id2b*-targeting MOs, we generated *pCS2+dlb-EGFP, pCS2+n3-EGFP* and *pCS2+id2b-EGFP* constructs that contained either *dlb, n3*, or *id2b* MO target site upstream of the EGFP initiator AUG. Plasmids were linearized with Not1 restriction enzyme and capped sense RNA was synthesized using SP6 RNA polymerase using Invitrogen’s mMESSAGE mMACHINE™ SP6 Transcription Kit (Invitrogen, #AM1340) according to manufactures directions. Following purification, the capped RNA was dissolved in nuclease free water containing 0.2% phenol red and injected with experimental or control MO into single cell zebrafish embryos. Each embryo received approximately 50 pg of RNA and 250 pg of control or experimental MO. MO’s were delivered to adult fish via intravitreal injection at the time of retinal injury and cellular uptake was facilitated by electroporation as previously described(Fausett et al 2008).

### Luciferase assay

A 3kb region upstream of the *hey1* gene’s transcription start site was amplified by PCR and cloned into Xho1/Kpn1 sites of the *pXP1* plasmid to create *3kb-hey1:luciferase*. Using Q5^®^ Site-Directed Mutagenesis Kit (NEB, E0554), we generated mutations in each of the two most highly conserved Rbpj binding sites (m1 and m2, Fig. 7a). Mutations were confirmed by DNA sequencing. To study *hey1* promoter activity, we transfected HEK 293T cells plated in 24 well tissue culture plates with wt and mutant *3kb-hey1:luciferase*, +/- CMV:NICD and *SV40:Renilla* as internal control using lipofectamine 3000 (Invitrogen). Two days after transfection, cells were washed with PBS, lysed, and luciferase and Renilla activities measured using a Luminometer (Turner Biosystems). Luciferase activity was normalized to Renilla activity.

### Heat shock and pharmacological inhibitors

For heat shock, fish were immersed in a water bath at 37°C for 1 hr before returning to system water at 28°C. For extended periods of heat shock, this was repeated every 6 hrs. To inhibit Notch signaling we used DAPT (Cayman, Cat # 13197) prepared in DMSO as a 10mM stock and diluted 1/250 in fish water for immersion. Control fish were treated with vehicle. Fish were exposed to DAPT for 24 hours.

### Immunofluorescence and TUNEL assay

Zebrafish samples were prepared for immunofluorescence as previously described(Fausett & Goldman 2006, Ramachandran et al 2010a, Ramachandran 2010). Primary antibodies used in this study: Rabbit anti-GFP, Thermo Fisher, Cat. # A6455 (1/1000); Mouse anti-mCherry, Abcam, Cat. # ab125096 (1:500); and mouse anti-glutamine synthetase (GS), Sigma-Aldrich, Cat. #MAB302 (1/500). Secondary antibodies used were: Alexa Flour 555 Donkey anti Mouse-IgG (H+L), Thermo Fisher Cat. # A31570 (1:500); Alexa flour 555 Donkey anti Rabbit IgG (H+L), Thermo Fisher, Cat # A31572 (1:500); Alexa Flour 488 donkey anti mouse Thermo Fisher Cat. # A21202 (1:500); Alexa Flour 488 goat anti rabbit Thermo Fisher Cat. # A11008 (1:500).

We used an *in situ* Cell Death Fluorescein Kit (Millipore Sigma, Cat. # 11684795910) to detect cells undergoing apoptosis.

### Microscopy and cell quantification

Images were captured by a Zeiss Axiophot fluorescence microscope or a Leica DM2500 microscope. EdU and BrdU labelling were used to identify and quantify proliferating cells in retinal sections as previously described(Fausett & Goldman 2006, Ramachandran et al 2010a, Wan & Goldman 2017, Wan et al 2012, Wan et al 2014). The width of the proliferative zone was quantified at the center of the injury site where the largest number of proliferating cells reside.

### Statistical analysis

Unless otherwise indicated, sample size is 3 fish and experiments were repeated at least 3 times. Statistical analyses were performed in GraphPad Prism. Error bars are standard deviation (SD). Twotailed Student’s *t* test was used for single parameter comparison; all measurements were taken from distinct samples.

### Data sharing

Raw data have been deposited in GEO under the superset accession number GSE160179 and will be released upon acceptance of manuscript for publication.

## Acknowledgments

This work was supported by grants from the NIH (NEI RO1 EY018132 and NEI RO1 EY027310) and the Gilbert Family Foundation Vision Restoration Initiative. We thank David Hyde, Notre Dame, for *gfap:GFP* transgenic fish; Michael Parsons, University California, Irvine, for *tp1:mCherry* transgenic fish; Mi-Sun Lee for generating *ubi:Cas9* transgenic fish; Elizabeth Mills for generating *u6:dll4-gRNAs_1,2_* and *u6:n3-gRNAs_1,2_* transgenic fish; Muchu Zhou and Zachary Rekowski for maintaining our zebrafish colony; and all members of the Goldman lab for their comments on this work. We also acknowledge and thank the University of Michigan’s Cell Sorting Core, Advanced Genomics Core, and Bioinformatics Core.

## Author contributions

AS designed and performed all experiments except ATACseq and generation of CRISPR gene edited fish. SD performed ATACseq experiments and generated CRISPR gene edited fish. JJ analyzed bulk RNAseq and scRNAseq data and helped in transfections. DG conceived, designed, and supervised the work and co-wrote the manuscript and helped generate transgenic fish. All authors reviewed and edited the manuscript.

## SUPPLEMENTARY FIGURE LEGENDS

**Figure S1:**
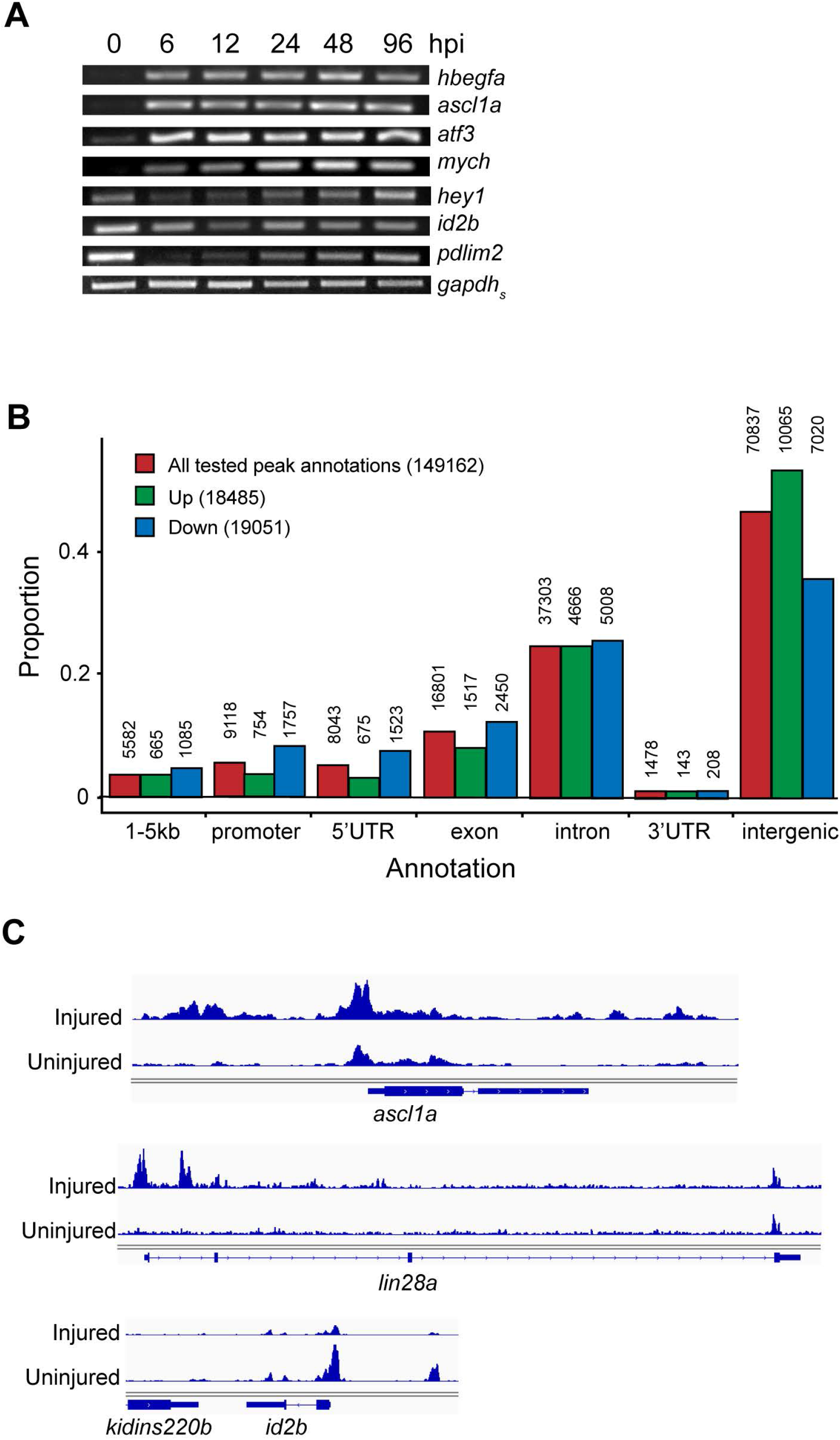
Injury-regulated genes and chromatin accessibility. **A**, PCR and agarose gel analysis of select regeneration-associated genes at various times post injury. **B**, Annotation of injury-regulated ATACseq data to different genic features. **C**, Select ATACseq genomic tracts generated from GFP+ MG FACS purified from uninjured and needle poke injured *gfap:GFP* retinas.

**Figure S2:**
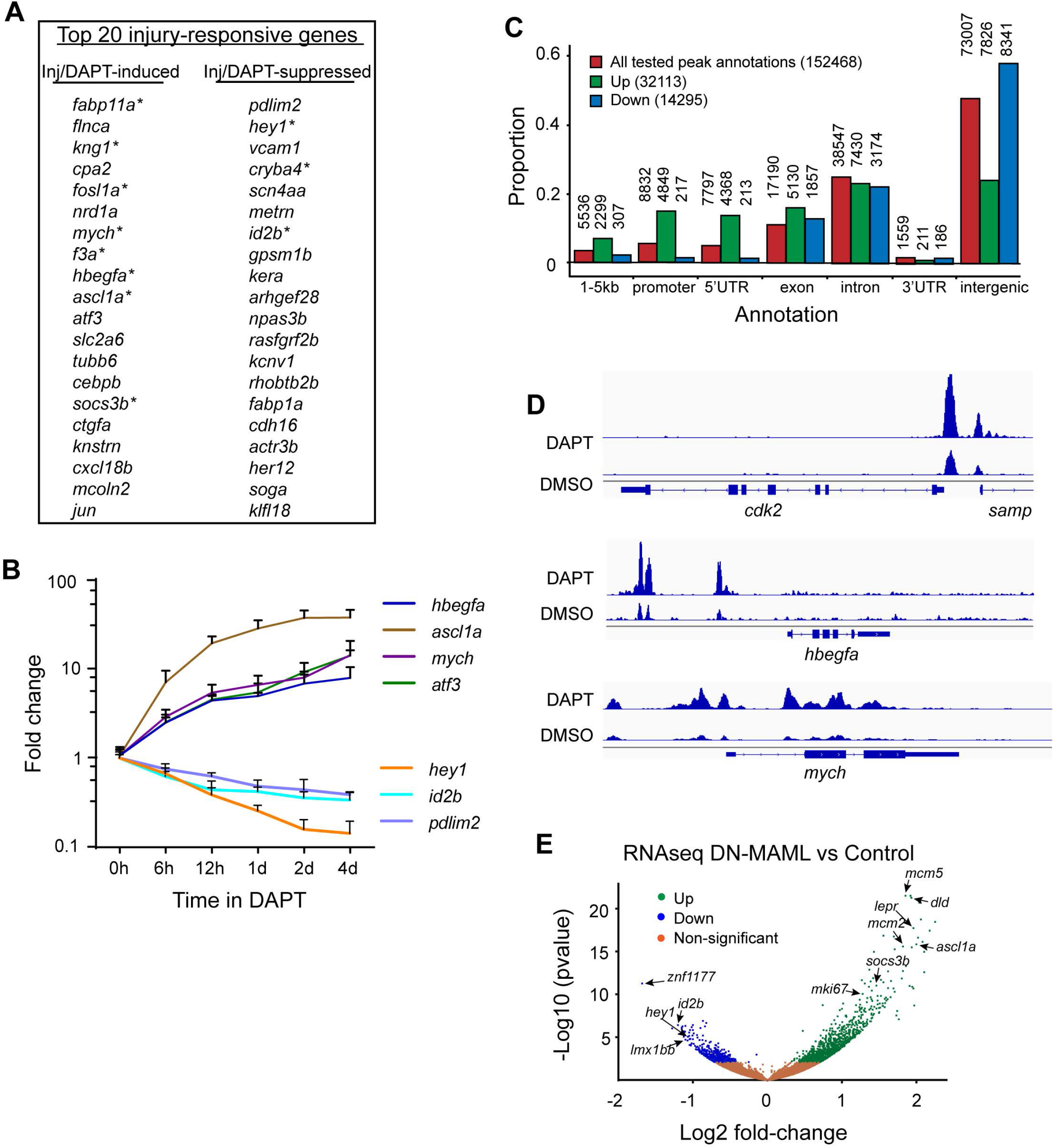
Notch-regulated genes and chromatin accessibility. **A**, Top 20 injury-responsive genes that are also regulated in a similar direction by DAPT-treatment. **B**, qPCR time course analysis of select DAPT-regulated genes in the uninjured retina. **C**, RNAseq FPKM values for select MG identity genes; \**P*=0.0385 (*gfap*). **D**, Annotation of DAPT-regulated ATACseq data to different genic features. **E**, Select ATACseq genomic tracts using GFP+ MG purified from uninjured *gfap:GFP* retinas treated +/-DAPT. **F**, Volcano plot of MG RNAseq data from *gfap:mCherry* and *gfap:mCherrry;hsp70:DN-MAML* heat shock treated fish.

**Figure S3:**
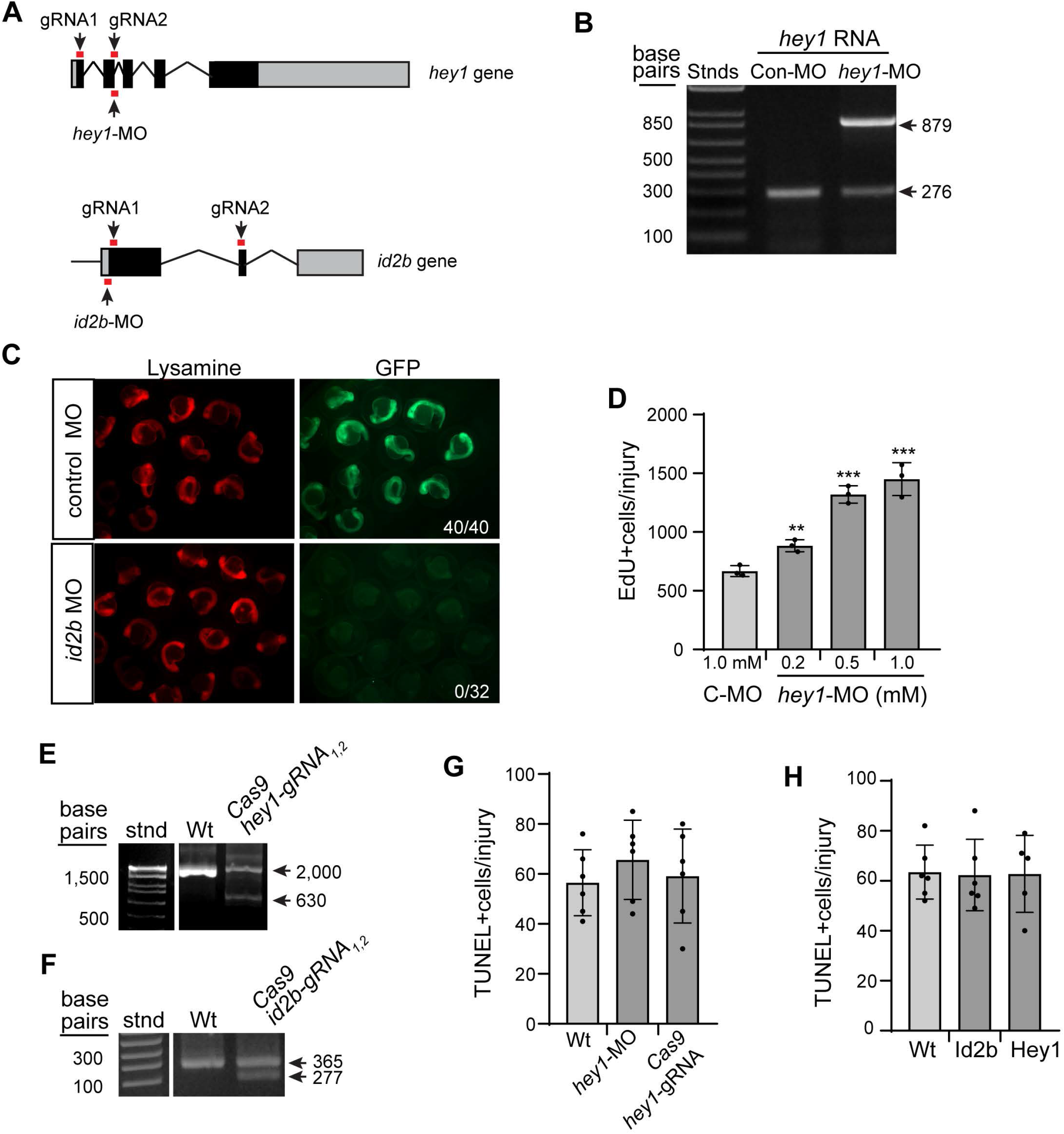
Hey1 and Id2b regulate retina regeneration. **A**, Illustration of the *hey1* gene with MO and gRNA target sites indicated. **B**, PCR analysis of *hey1*-MO treatment stimulating *hey1* intron retention. **C**, *id2b*-MO suppressed *id2b*-GFP chimeric RNA expression in zebrafish embryos. Numbers in panels refer to number of lissamine+ fish with GFP+ phenotype. **D**, Quantification of EdU+ cells using Click-iT chemistry in *hey1*-MO treated retina; \*\**P*=0.0054 (0.2mM); \*\*\**P*=0.0002 (0.5mM); \*\*\**P*=0.0008 (1.0mM). **E**, **F**, Single cell embryos were injected with Cas9 RNA and the indicated gRNAs and raised to adults. Shown is PCR analysis of genomic DNA from fin clips of adult fish. **G**, **H**, Quantification of TUNEL+ cells in the injured retina.

**Figure S4:**
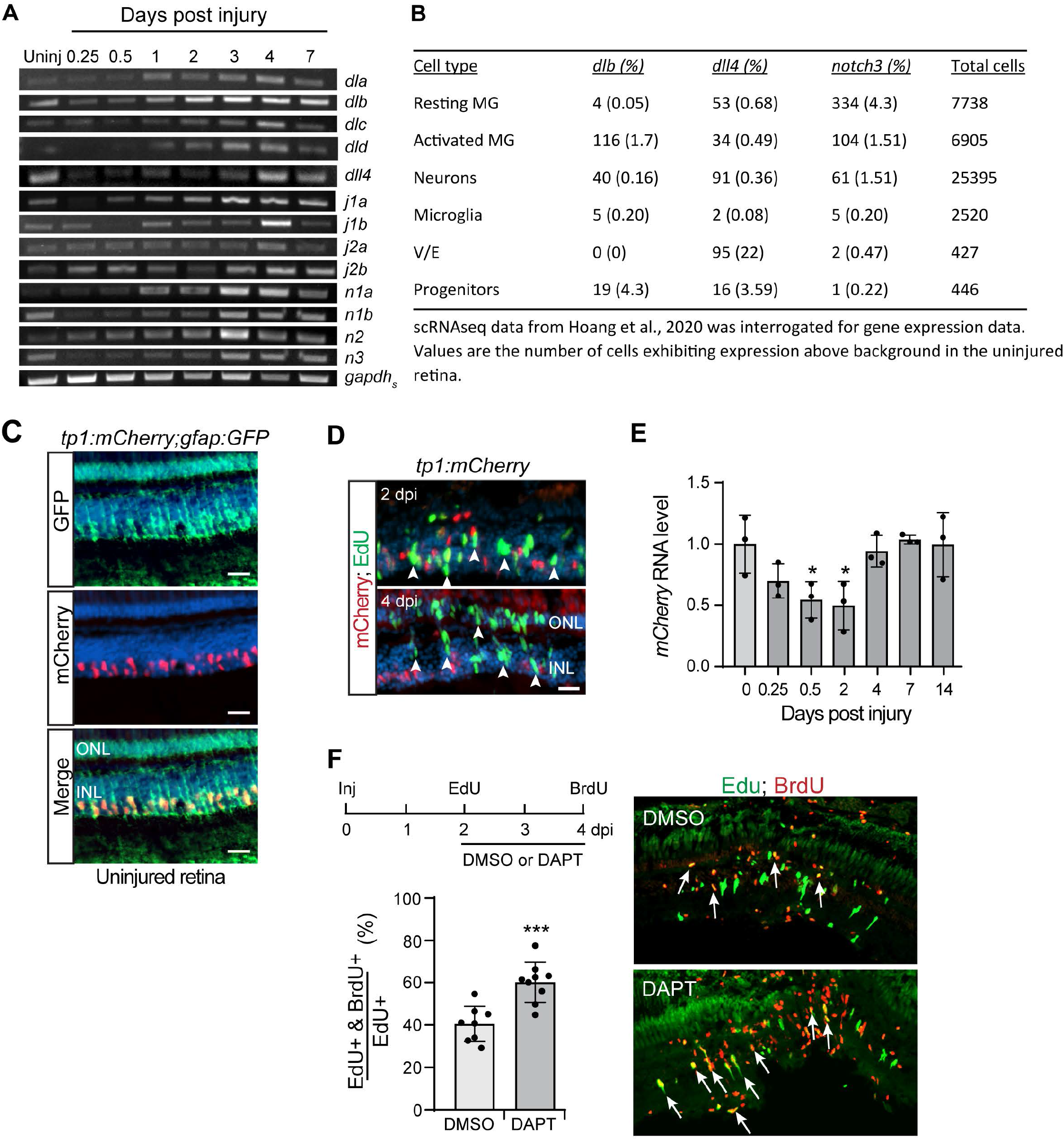
Injury-dependent changes in Notch ligand and receptor gene expression. **A**, Total retinal RNA was used for PCR analysis of Notch ligand and receptor gene expression at various times post injury. **B**, Percentage of indicated retinal cell types expressing the indicated gene based on data from Hoang et al., 2020. **C**, GFP and mCherry immunofluorescence on adult retinal sections from *tp1:mCherry;gfap:GFP* transgenic fish. **D**, Representative fluorescent image of mCherry immunofluorescence (red) and EdU Click-iT chemistry (green) on retinal sections at 2 and 4 dpi. Arrows point to EdU+/mCherry-MG-derived progenitors. **E**, Graph is qPCR quantification of *mCherry* RNA expression in adult retina of *tp1:mCherry* fish at various times post needle poke injury. Compared to 0 dpi, \**P*=0.0478 at 0.5 and 2 days post injury. **F**, Top: Experimental time line. Graph: Percentage of EdU+ cells that are also BrdU+. Right: Representative fluorescent images following DAPT treatment from 2-4 dpi in the needle pole injured retina (EdU+ cells are green and BrdU+ cells are red). Arrows point to double labelled cells. Abbreviations: ONL, outer nuclear layer; INL, inner nuclear layer; GCL, ganglion cell layer. Scale bar is 100 μm.

**Figure S5:**
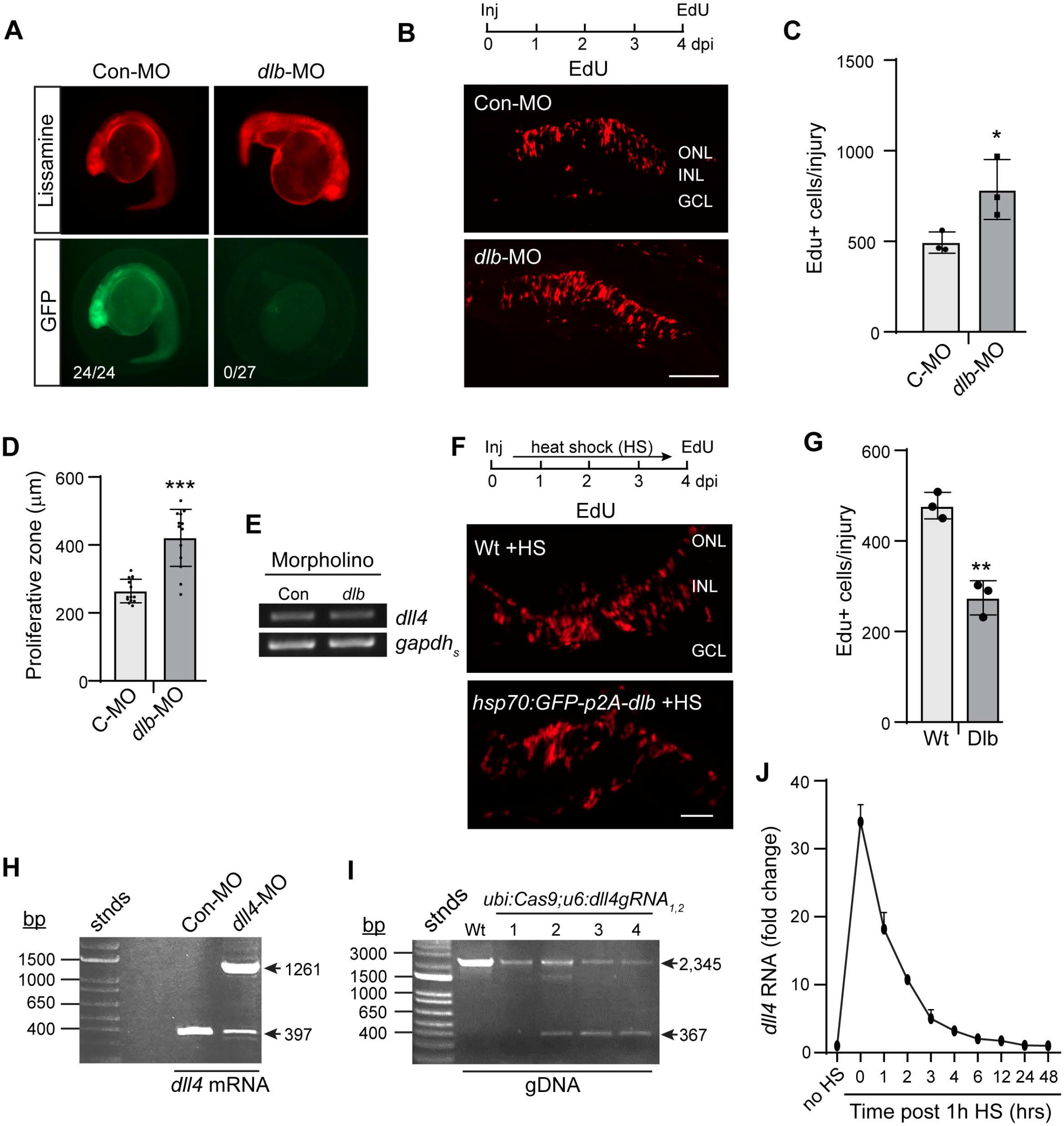
Notch ligand encoding genes that regulate MG’s injury response. **A**, *dlb*-MO suppresses *dlb*-GFP chimeric RNA expression in embryos. Numbers in panels refer to number of lissamine+ fish with GFP+ phenotype. **B**, Experimental time line and EdU Click-iT chemistry on retinal sections to detect proliferating cells. **C**, Quantification of EdU+ cells in control and Dlb knockdown retinas; \**P*<0.0446. **D**, Quantification of the proliferative zone in the injured zebrafish retina treated with control and *dlb*-MO; \*\*\**P*<0.0001. **E**, PCR analysis of *dll4* RNA expression in retinas treated with control and *dlb*-MO. **F**, Experimental time line and EdU Click-iT chemistry on retinal sections to identify proliferating cells. **G**, Quantification of EdU+ cells in control (Wt) and Dlb overexpressing (Dlb) transgenic fish; \*\**P*=0.0018. **H**, PCR shows *dll4*-MO stimulates intron retention in the *dll4* RNA. **I**, CRISPR-based gene editing detected in genomic DNA from *ubi:Cas9;u6:dll4gRNA_1,2_* transgenic fish. **J**, qPCR analysis of *dll4* transgene induction in *hsp70:dll4-p2A-GFP* fish after heat shock. Abbreviations: MO, morpholino; C-MO or Con-MO, control-MO; n3-MO, notch3-MO; ONL, outer nuclear layer; INL, inner nuclear layer; GCL, ganglion cell layer. Scale bars are 100 μm.

**Figure S6:**
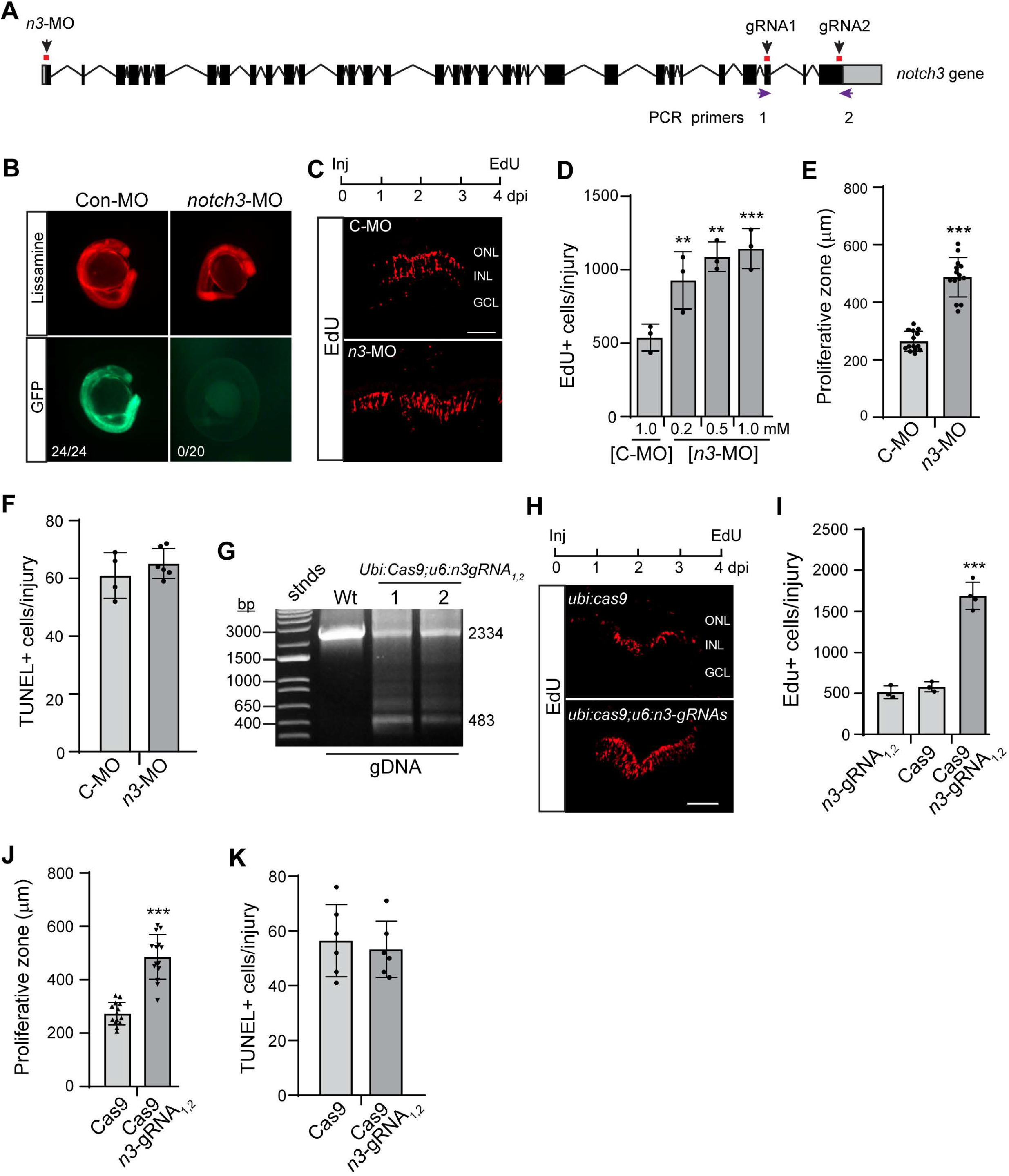
Notch3 regulates MG’s threshold response to injury. **A**, Diagram of the *notch3* gene with MO and gRNA target sites indicated. **B**, *notch3*-MO suppresses *notch3-GFP* chimeric RNA expression in embryos. Numbers in panels refer to number of lissamine+ fish with GFP+ phenotype. **C**, **H**, EdU Click-iT assay on retinal sections at 4 dpi to detect proliferating cells. **D**, **I**, Quantification of the number of EdU+ cells at 4 dpi; (**D**) Compared to C-MO, \*\**P*=0.0019 (0.2mM), \*\**P*=0.0015 (0.5mM), \*\*\**P*=0.0006 (1mM); (**I**) compared to Cas9 \*\*\**P*=0.0002. **E**, **J**, Quantification of the width of the proliferative zone in retinas at 4 dpi; (**E**) \*\*\**P*<0.0001; (**J**) \*\*\**P*<0.0001; **F**, **K**, Quantification of TUNEL+ cells at 1 dpi. **G**, CRISPR-based gene editing detected by PCR using genomic DNA from transgenic fish. Abbreviations: MO, morpholino; C-MO or Con-MO, control-MO; n3-MO, notch3-MO; ONL, outer nuclear layer; INL, inner nuclear layer; GCL, ganglion cell layer. Scale bars are 100 μm.

## Supplementary Tables

**Table S1:** Inj reg genes & chromatin. Related to Figure 1.

**Table S2:** Inj reg ATAC genes, promoters, & 1-5kb regions. Related to Figure 1.

**Table S3:** Inj and DAPT coordinately regulated genes. Related to Figure 2.

**Table S4**: DAPT & inj regulated ATACseq. Related to Figure 2.

**Table S5**: DAPT & DN-MAML intersection with regeneration-associated genes. Related to Figure 2.

**Table S6:** PCR heat map data. Related to Figure 4.

**Table S7:** Primer, MO, gRNA list. Related to Material and methods.

## References

Alunni A, Krecsmarik M, Bosco A, Galant S, Pan L, et al. 2013. Notch3 signaling gates cell cycle entry and limits neural stem cell amplification in the adult pallium. Development 140: 3335–47

Alvarez Y, Cederlund ML, Cottell DC, Bill BR, Ekker SC, et al. 2007. Genetic determinants of hyaloid and retinal vasculature in zebrafish. BMC Dev Biol 7: 114

Attwell D, Buchan AM, Charpak S, Lauritzen M, Macvicar BA, Newman EA. 2010. Glial and neuronal control of brain blood flow. Nature 468: 232–43

Bernardos RL, Lentz SI, Wolfe MS, Raymond PA. 2005. Notch-Delta signaling is required for spatial patterning and Muller glia differentiation in the zebrafish retina. Dev Biol 278: 381–95

Buenrostro JD, Wu B, Chang HY, Greenleaf WJ. 2015. ATAC-seq: A Method for Assaying Chromatin Accessibility Genome-Wide. Curr Protoc Mol Biol 109: 21.29.1–21.29.9

Campbell LJ, Hobgood JS, Jia M, Boyd P, Hipp RI, Hyde DR. 2021. Notch3 and DeltaB maintain Muller glia quiescence and act as negative regulators of regeneration in the light-damaged zebrafish retina. Glia 69: 546–66

Conner C, Ackerman KM, Lahne M, Hobgood JS, Hyde DR. 2014. Repressing notch signaling and expressing TNFalpha are sufficient to mimic retinal regeneration by inducing Muller glial proliferation to generate committed progenitor cells. J Neurocsci 34: 14403–19

Elsaeidi F, Macpherson P, Mills EA, Jui J, Flannery JG, Goldman D. 2018. Notch Suppression Collaborates with Ascl1 and Lin28 to Unleash a Regenerative Response in Fish Retina, But Not in Mice. J Neurosci 38: 2246–61

Engler A, Zhang R, Taylor V. 2018. Notch and Neurogenesis. Adv Exp Med Biol 1066: 223–34

Fausett BV, Goldman D. 2006. A role for alpha1 tubulin-expressing Muller glia in regeneration of the injured zebrafish retina. J Neurosci 26: 6303–13

Fausett BV, Gumerson JD, Goldman D. 2008. The proneural basic helix-loop-helix gene ascl1a is required for retina regeneration. J Neurosci 28: 1109–17

Garhofer G, Chua J, Tan B, Wong D, Schmidl D, Schmetterer L. 2020. Retinal Neurovascular Coupling in Diabetes. J Clin Med 9: 2829

Goldman D. 2014. Muller glial cell reprogramming and retina regeneration. Nature reviews. Neuroscience 15: 431–42

Gorsuch RA, Lahne M, Yarka CE, Petravick ME, Li J, Hyde DR. 2017. Sox2 regulates Muller glia reprogramming and proliferation in the regenerating zebrafish retina via Lin28 and Ascl1a. Exp Eye Res 161: 174–92

Hoang T, Wang J, Boyd P, Wang F, Santiago C, et al. 2020. Gene regulatory networks controlling vertebrate retinal regeneration. Science 370: eabb8598

Hwang WY, Fu Y, Reyon D, Maeder ML, Tsai SQ, et al. 2013. Efficient genome editing in zebrafish using a CRISPR-Cas system. Nat Biotechnol 31: 227–9

Imayoshi I, Sakamoto M, Yamaguchi M, Mori K, Kageyama R. 2010. Essential roles of Notch signaling in maintenance of neural stem cells in developing and adult brains. J Neurosci 30: 3489–98

Iribarne M, Hyde DR, Masai I. 2019. TNFalpha Induces Muller Glia to Transition From Non-proliferative Gliosis to a Regenerative Response in Mutant Zebrafish Presenting Chronic Photoreceptor Degeneration. Front Cell Dev Biol 7: 296

Ivanova E, Alam NM, Prusky GT, Sagdullaev BT. 2019. Blood-retina barrier failure and vision loss in neuron-specific degeneration. JCI Insight 4: e126747

Jadhav AP, Cho SH, Cepko CL. 2006. Notch activity permits retinal cells to progress through multiple progenitor states and acquire a stem cell property. Proc Natl Acad Sci U S A 103: 18998–9003

Johnson K, Barragan J, Bashiruddin S, Smith CJ, Tyrrell C, et al. 2016. Gfap-positive radial glial cells are an essential progenitor population for later-born neurons and glia in the zebrafish spinal cord. Glia 64: 1170–89

Kassen SC, Ramanan V, Montgomery JE, C TB, Liu CG, et al. 2007. Time course analysis of gene expression during light-induced photoreceptor cell death and regeneration in albino zebrafish. Dev Neurobiol 67: 1009–31

Kawai H, Kawaguchi D, Kuebrich BD, Kitamoto T, Yamaguchi M, et al. 2017. Area-Specific Regulation of Quiescent Neural Stem Cells by Notch3 in the Adult Mouse Subependymal Zone. J Neurosci 37: 11867–80

Kim EJ, Leung CT, Reed RR, Johnson JE. 2007. In vivo analysis of Ascl1 defined progenitors reveals distinct developmental dynamics during adult neurogenesis and gliogenesis. J Neurosci 27: 12764–74

Lahne M, Nagashima M, Hyde DR, Hitchcock PF. 2020. Reprogramming Muller Glia to Regenerate Retinal Neurons. Annu Rev Vis Sci 6: 171–93

Lee MS, Wan J, Goldman D. 2020. Tgfb3 collaborates with PP2A and notch signaling pathways to inhibit retina regeneration. Elife 9: e55137

Lenkowski JR, Raymond PA. 2014. Muller glia: Stem cells for generation and regeneration of retinal neurons in teleost fish. Progress in retinal and eye research 40: 94–123

Leslie JD, Ariza-McNaughton L, Bermange AL, McAdow R, Johnson SL, Lewis J. 2007. Endothelial signalling by the Notch ligand Delta-like 4 restricts angiogenesis. Development 134: 839–44

Lessieur EM, Song P, Nivar GC, Piccillo EM, Fogerty J, et al. 2019. Ciliary genes arl13b, ahi1 and cc2d2a differentially modify expression of visual acuity phenotypes but do not enhance retinal degeneration due to mutation of cep290 in zebrafish. PLoS One 14: e0213960

Loffler K, Schafer P, Volkner M, Holdt T, Karl MO. 2015. Age-dependent Muller glia neurogenic competence in the mouse retina. Glia 63: 1809–24

MacDonald RB, Randlett O, Oswald J, Yoshimatsu T, Franze K, Harris WA. 2015. Muller glia provide essential tensile strength to the developing retina. J Cell Biol 210: 1075–83

Maillard I, Tu L, Sambandam A, Yashiro-Ohtani Y, Millholland J, et al. 2006. The requirement for Notch signaling at the beta-selection checkpoint in vivo is absolute and independent of the pre-T cell receptor. J Exp Med 203: 2239–45

Maillard I, Weng AP, Carpenter AC, Rodriguez CG, Sai H, et al. 2004. Mastermind critically regulates Notch-mediated lymphoid cell fate decisions. Blood 104: 1696–702

Montgomery JE, Parsons MJ, Hyde DR. 2010. A novel model of retinal ablation demonstrates that the extent of rod cell death regulates the origin of the regenerated zebrafish rod photoreceptors. J Comp Neurol 518: 800–14

Moreno-Mateos MA, Vejnar CE, Beaudoin JD, Fernandez JP, Mis EK, et al. 2015. CRISPRscan: designing highly efficient sgRNAs for CRISPR-Cas9 targeting in vivo. Nat Methods 12: 982–8

Nagashima M, Barthel LK, Raymond PA. 2013. A self-renewing division of zebrafish Muller glial cells generates neuronal progenitors that require N-cadherin to regenerate retinal neurons. Development 140: 4510–21

Nam Y, Sliz P, Song L, Aster JC, Blacklow SC. 2006. Structural basis for cooperativity in recruitment of MAML coactivators to Notch transcription complexes. Cell 124: 973–83

Nelson BR, Ueki Y, Reardon S, Karl MO, Georgi S, et al. 2011. Genome-wide analysis of Muller glial differentiation reveals a requirement for Notch signaling in postmitotic cells to maintain the glial fate. PLoS One 6: e22817

Parsons MJ, Pisharath H, Yusuff S, Moore JC, Siekmann AF, et al. 2009. Notch-responsive cells initiate the secondary transition in larval zebrafish pancreas. Mech Dev 126: 898–912

Powell C, Grant AR, Cornblath E, Goldman D. 2013. Analysis of DNA methylation reveals a partial reprogramming of the Muller glia genome during retina regeneration. Proc Natl Acad Sci U S A 110: 19814–9

Ramachandran R, Fausett BV, Goldman D. 2010a. Ascl1a regulates Muller glia dedifferentiation and retinal regeneration through a Lin-28-dependent, let-7 microRNA signalling pathway. Nat Cell Biol 12: 1101–7

Ramachandran R, Reifler, A., Parent, J.M and Goldman, D. 2010. Conditional gene expression and lineage tracing of tuba1a expressing cells during zebrafish development and retina regeneration. J Comp Neurol 518: 4196–212

Ramachandran R, Zhao XF, Goldman D. 2011. Ascl1a/Dkk/{beta}-catenin signaling pathway is necessary and glycogen synthase kinase-3{beta} inhibition is sufficient for zebrafish retina regeneration. Proc Natl Acad Sci U S A 108: 15858–63

Ramachandran R, Zhao XF, Goldman D. 2012. Insm1a-mediated gene repression is essential for the formation and differentiation of Muller glia-derived progenitors in the injured retina. Nat Cell Biol 14: 1013–23

Reichenbach A, Bringmann A. 2013. New functions of Muller cells. Glia 61: 651–78

Riesenberg AN, Conley KW, Le TT, Brown NL. 2018. Separate and coincident expression of Hes1 and Hes5 in the developing mouse eye. Dev Dyn 247: 212–21

Sakamoto M, Hirata H, Ohtsuka T, Bessho Y, Kageyama R. 2003. The basic helix-loop-helix genes Hesr1/Hey1 and Hesr2/Hey2 regulate maintenance of neural precursor cells in the brain. J Biol Chem 278: 44808–15

Scheer N, Groth A, Hans S, Campos-Ortega JA. 2001. An instructive function for Notch in promoting gliogenesis in the zebrafish retina. Development 128: 1099–107

Sentmanat MF, Peters ST, Florian CP, Connelly JP, Pruett-Miller SM. 2018. A Survey of Validation Strategies for CRISPR-Cas9 Editing. Sci Rep 8: 888

Sifuentes CJ, Kim JW, Swaroop A, Raymond PA. 2016. Rapid, Dynamic Activation of Muller Glial Stem Cell Responses in Zebrafish. Invest Ophthalmol Vis Sci 57: 5148–60

Than-Trong E, Ortica-Gatti S, Mella S, Nepal C, Alunni A, Bally-Cuif L. 2018. Neural stem cell quiescence and stemness are molecularly distinct outputs of the Notch3 signalling cascade in the vertebrate adult brain. Development 145: dev161034

Vejnar CE, Moreno-Mateos MA, Cifuentes D, Bazzini AA, Giraldez AJ. 2016. Optimized CRISPR-Cas9 System for Genome Editing in Zebrafish. Cold Spring Harb Protoc 2016: doi:10.1101/pdb.prot086850

Wan J, Goldman D. 2016. Retina regeneration in zebrafish. Curr Opin Genet Dev 40: 41–47

Wan J, Goldman D. 2017. Opposing Actions of Fgf8a on Notch Signaling Distinguish Two Muller Glial Cell Populations that Contribute to Retina Growth and Regeneration. Cell Rep 19: 849–62

Wan J, Ramachandran R, Goldman D. 2012. HB-EGF is necessary and sufficient for Muller glia dedifferentiation and retina regeneration. Dev Cell 22: 334–47

Wan J, Zhao XF, Vojtek A, Goldman D. 2014. Retinal Injury, Growth Factors, and Cytokines Converge on beta-Catenin and pStat3 Signaling to Stimulate Retina Regeneration. Cell Reports 9: 285–97

Wapinski OL, Lee QY, Chen AC, Li R, Corces MR, et al. 2017. Rapid Chromatin Switch in the Direct Reprogramming of Fibroblasts to Neurons. Cell Rep 20: 3236–47

Xia W. 2019. gamma-Secretase and its modulators: Twenty years and beyond. Neurosci Lett 701: 162–69

Yin L, Maddison LA, Li M, Kara N, LaFave MC, et al. 2015. Multiplex Conditional Mutagenesis Using Transgenic Expression of Cas9 and sgRNAs. Genetics 200: 431–41

Zaucker A, Mercurio S, Sternheim N, Talbot WS, Marlow FL. 2013. notch3 is essential for oligodendrocyte development and vascular integrity in zebrafish. Dis Model Mech 6: 1246–59

Zhao L, Borikova AL, Ben-Yair R, Guner-Ataman B, MacRae CA, et al. 2014a. Notch signaling regulates cardiomyocyte proliferation during zebrafish heart regeneration. Proc Natl Acad Sci U S A 111: 1403–8

Zhao XF, Wan J, Powell C, Ramachandran R, Myers MG, Jr., Goldman D. 2014b. Leptin and IL-6 Family Cytokines Synergize to Stimulate Muller Glia Reprogramming and Retina Regeneration. Cell Reports 9: 272–84

